# A multipurpose microhaplotype panel for genetic analysis of California Chinook salmon

**DOI:** 10.1101/2024.12.31.630931

**Authors:** Eric C. Anderson, Anthony J. Clemento, Matthew A. Campell, Devon E. Pearse, Anne K. Beulke, Cassie Columbus, Ellen Campbell, Neil F. Thompson, John Carlos Garza

## Abstract

Genetic methods have become an essential component of ecological investigation and conservation planning for fish and wildlife. Among these methods is the use of genetic marker data to identify individuals to populations, or stocks, of origin. More recently, methods that involve genetic pedigree reconstruction to identify relationships between individuals within populations have also become common. We present, here, a novel set of multi-allelic microhaplotype genetic markers for Chinook salmon which provide unprecedented resolution for population discrimination and relationship identification from a rapidly and economically assayed panel of markers. We show how this set of microhaplotypes provides definitive power to identify all known lineages of Chinook salmon in California. The inclusion of genetic loci that have known associations with phenotype and that were identified as outliers in examination of whole genome sequence data, allows resolution of stocks that are not highly genetically differentiated but are phenotypically distinct and managed as such. This same set of multiallelic genetic markers have ample variation to accurately identify parent-offspring and full-sibling pairs in all California populations, including the genetically depauperate winter-run lineage. Validation of this marker panel in coastal salmon populations not previously studied with modern genetic methods, also reveals novel biological insights, including the presence of a single copy of a haplotype for a phenotype that has not been documented in that part of the species range, and a clear signal of mixed ancestry for a salmon population that is on the geographic margins of the primary evolutionary lineages present in California.

## Introduction

Genetic markers have long been used in the management of Pacific salmon, and salmon conservation and management have been at the forefront of a number of advances in molecular ecology, from data generation techniques (Clemento *et al*., 2011; Campbell *et al*., 2015; McKinney *et al*., 2017; Baetscher *et al*., 2018) to statistical methodology (Smouse *et al*., 1990; Anderson & Thompson, 2002; Pella & Masuda, 2006; Anderson, 2010). Two broadly applicable techniques that have been actively fostered by the Pacific salmon research community are genetic stock identification (GSI: Milner *et al*., 1982; Beacham *et al*., 2004; Seeb *et al*., 2007) and parentage based tagging (PBT: Anderson & Garza, 2006; Garza & Anderson, 2007; Abadía-Cardoso *et al*., 2013; Steele *et al*., 2013).

In the 1980s, electrophoretically detectable genetic variation, in the form of allozymes (Ayala & Powell, 1972; Allendorf & Phelps, 1981), was used to establish a program of GSI for Chinook salmon, *Oncorhynchus tshawytscha* (Milner *et al*.,1982). Extensive sampling revealed that these allozyme markers exhibited different allele frequencies among major lineages, or stocks, of Chinook salmon on the West Coast. These allele frequency differences make GSI possible, and Milner *et al*. (1985) soon showed that proportions of stocks in the Washington state coastal troll fishery could be estimated by GSI. Since that time, with the development of novel molecular markers, and, now with increasing capacity to sequence genomic material, the scope and scale of GSI has expanded considerably.

By using greater numbers of more variable markers than the allozymes available in the 1980s it is possible to accurately identify the population of origin of individual fish, rather than simply estimating aggregated stock proportions. It is also possible to resolve populations of fish that are much more closely related than before. Furthermore, reference data sets with genotypes from hundreds of populations throughout the range of multiple species of salmon and other anadromous species (Seeb *et al*., 2007; Gilbey *et al*., 2018; Barclay &Habicht, 2019) now exist, and are routinely used to assign fish caught thousands of kilometers from their natal streams to their stock of origin. Applications include estimating fishery composition (Satterthwaite *et al*., 2015), providing real-time information for genetics-informed fishery closures (Beacham *et al*., 2004), assessing the spatial distribution of different stocks in the ocean (Urawa *et al*., 2009) and their temporal distribution in upstream migrations (Hess *et al*., 2014), and monitoring bycatch (Hasselman *et al*., 2016) or illegal captures (Wilmot *et al*., 1999) in marine fisheries.

Although sequencing costs continue to decline, they are high enough that there remains a tradeoff between reference baselines that include information from a large number of populations across a broad scale, and those that have been tailored to distinguish between closely related populations on a smaller, regional scale. Because of cost considerations, reference baselines that include populations across a broad spatial scale may include only a few populations from each sub-region. Furthermore, baselines tailored to a specific region often assemble markers that show allele frequency differences between the closely related—and hence difficult to resolve—stocks within the region. Consequently, regionally targeted baselines typically outperform broad-scale baselines in resolving populations within the region.

Over the last decade an additional genetic method, parentage based tagging, or PBT, (Anderson & Garza, 2005; Steele *et al*., 2019) has become established as an extremely valuable management tool for Pacific salmon. The availability of such family-based methods adds another factor to consider when developing a GSI baseline. Since the first proposal (Anderson & Garza, 2005) to replace or augment the coded-wire tag programme (Nandor *et al*., 2010) with PBT, it has been noted that one of the major advantages of a genetic program for PBT is that the genetic markers used for PBT could also be useful for GSI (and vice versa). Thus, any panel of markers to be used for GSI (or PBT) should also be evaluated on its utility for PBT (or GSI).

PBT has been remarkably successful in fisheries management, having been used for over a decade in the management of Chinook salmon and steelhead trout in major basins of the Columbia River (Steele *et al*., 2019; Horn *et al*., 2023), and having been employed to dramatically further our understanding of the genetic inheritance of key traits in salmonids (Abadía-Cardoso *et al*., 2013; Beulke *et al*., 2023). However, PBT is just one subset of a whole family of statistical genetic methods employing relationship inference to learn about populations. For example, inference of the full siblings amongst a sample of fish provides information about the effective number of adults producing offspring (Waples & Waples, 2011; Wang, 2023), which can be a valuable source of information when sampling of the adults is not possible. Accordingly, marker panels should also be evaluated on their capacity to resolve full-sibling relationships.

Chinook salmon are the largest of the Pacific salmonids and have historically been the target of extremely high-value and culturally important fisheries (Myers *et al*.,1998). Moreover, because of their high degree of ecotypic variation, they have provided fishery opportunities in many different seasons and geographic locations (Healey, 1991). However, recent declines in population sizes, from the Yukon River in the Arctic north, to the southern extent of their range in California, has led to multiple fishery closures to protect less productive stocks (Lindley *et al*., 2009). The co-occurrence of fish from relatively productive and relict populations reaches its paragon in California, with the largest remaining ocean fisheries for Chinook salmon targeting the Central Valley fall-run stock that spawns in different tributaries of the same river basin as the highly endangered and phenotypically distinct winter-run stock and the threatened spring-run stocks (Satterthwaite *et al*., 2015).

Here, we present a reference baseline for Chinook salmon, focusing on the regional scale of rivers within the state of California, and particularly targeted to the complex population structure of Chinook salmon within the California Central Valley (CCV). Chinook salmon of the two main CCV river basins—the Sacramento and the San Joaquin—exhibit the greatest run-timing diversity within the species. With four recognized ecotypes, delineated primarily on the basis of run timing (fall-, late-fall-, winter-, and spring-run), adult Chinook salmon can be found migrating or residing in freshwater every month of the year in California (Fisher, 1994).

In spite of their distinct phenotypes and life history patterns, the four ecotypes of Chinook salmon in the Central Valley are closely related (Clemento *et al*., 2014) and share recent common ancestry that is independent of ongoing migration between these lineages. As such, GSI has been particularly challenging in the CCV, with at least one of the distinct ecotypes, the late-fall-run, unresolvable with previous GSI marker sets and with unsatisfactory power for discriminating the protected spring-run stocks from the harvested fall-run stock (Seeb *et al*., 2007; Clemento *et al*., 2014).

In the following, we present a new reference baseline that includes 1,636 Chinook salmon individuals from 17 collections, genotyped at 204 loci, mostly microhaplotypes (Baetscher *et al*., 2018), from throughout the genome. This baseline is highly effective for GSI within California and the markers within it provide ample power for PBT, thus enabling a highly effective and efficient integrated GSI/PBT monitoring and evaluation program (Garza & Anderson, 2007; Beacham *et al*., 2021). We describe how we identified and compiled the markers, including several new markers that are particularly effective at distinguishing between closely related groups of fish, such as late-fall and fall-run Chinook salmon in the CCV. We also provide a comprehensive analysis of this set of markers for inference of parents and siblings within different groups of populations.

## Methods

### Population Sampling

We compiled samples of fish for the reference baselines from 13 locations within California, and one within Oregon. This set of populations and stocks includes all of the previously described lineages of Chinook salmon in the southern portion of the range and most of the populations that are known to have significant genetic differentiation in this region. These locations included sites in the two main tributaries (the Sacramento River and the San Joaquin River) within California’s Central Valley, and the two main tributaries (Klamath and Trinity Rivers) in the Klamath basin in Northern California, and three rivers of the California coast (Russian, Eel, and Smith) (Figure 1, Table 1).

**Table 1.** Collections of fish in the reference baseline. The Sampling Months column shows a pictorial representation of the proportion of each collection sampled in each month. BCkF were sampled as juveniles, all other collections were of adults. Colors in this column correspond to run-timing groups as described in the caption of Figure 1.

| Mainstem River<sup>1</sup> | Code | Location Name<sup>2</sup> | Run | Reporting Unit<sup>3</sup> | ESU<sup>4</sup> | N | N<sub>day</sub><sup>5</sup> | Sampling Months | Year Range |
| --- | --- | --- | --- | --- | --- | --- | --- | --- | --- |
| Rogue R. | CRHS | Cole Rivers H. | Spring | SO-NCal-Coast | SONCC | 50 | 0 | J F M A M J J A S O N D | 2004–2010 |
| Smith R. | SmRF | Smith R. | Fall | SO-NCal-Coast | SONCC | 46 | 46 | J F M A M J J A S O N D | 2010–2011 |
| Klamath R. | BCKf | Blue Ck. | Fall | SO-NCal-Coast | SONCC | 56 | 56 | J F M A M J J A S O N D | 2008–2008 |
| Klamath R. | IGHF | Iron Gate H. | Fall | Klamath-Trinity | UKTR | 93 | 92 | J F M A M J J A S O N D | 2017–2017 |
| Trinity R. | TRS | Trinity R. H. | Spring | Klamath-Trinity | UKTR | 46 | 46 | J F M A M J J A S O N D | 2012–2012 |
| Trinity R. | TRF | Trinity R. H. | Fall | Klamath-Trinity | UKTR | 41 | 41 | J F M A M J J A S O N D | 2012–2012 |
| Eel R. | ERF | Eel R. | Fall | Cent-Cal-Coast | CCC | 93 | 0 | J F M A M J J A S O N D | 2014–2014 |
| Russian R. | RRF | Warm Springs H. | Fall | Cent-Cal-Coast | CCC | 92 | 92 | J F M A M J J A S O N D | 2012–2012 |
| Sacramento R. | SRW | Livingston Stone NFH | Winter | CV-Winter | CVWR | 111 | 20 | J F M A M J J A S O N D | 2013–2014 |
| Sacramento R. | BCS | Butte Ck. | Spring | CV-Spring | CVSR | 116 | 116 | J F M A M J J A S O N D | 2004–2017 |
| Sacramento R. | MDS | Deer Ck. | Spring | CV-Spring | CVSR | 34 | 34 | J F M A M J J A S O N D | 2002–2002 |
| Sacramento R. | MDS | Mill Ck. | Spring | CV-Spring | CVSR | 50 | 50 | J F M A M J J A S O N D | 2002–2004 |
| Sacramento R. | FRHS | Feather R. H. | Spring | CV-Fall | CVSR | 202 | 202 | J F M A M J J A S O N D | 2010–2019 |
| Sacramento R. | FRHF | Feather R. H. | Fall | CV-Fall | CVFR | 106 | 106 | J F M A M J J A S O N D | 2008–2010 |
| Sacramento R. | BCF | Butte Ck. | Fall | CV-Fall | CVFR | 45 | 45 | J F M A M J J A S O N D | 2004–2016 |
| Sacramento R. | MDF | Deer Ck. | Fall | CV-Fall | CVFR | 24 | 24 | J F M A M J J A S O N D | 2002–2004 |
| Sacramento R. | MDF | Mill Ck. | Fall | CV-Fall | CVFR | 49 | 49 | J F M A M J J A S O N D | 2002–2004 |
| San Joaquin R. | SJRF | San Joaquin R. | Fall | CV-Fall | CVFR | 79 | 79 | J F M A M J J A S O N D | 2016–2016 |
| Sacramento R. | CHLF | Coleman NFH | Late-fall | CV-Late-Fall | CVFR | 261 | 261 | J F M A M J J A S O N D | 2011–2015 |

<sup>1</sup>R. = River.
<sup>2</sup>Ck. = Creek; NFH = National Fish Hatchery; R. = River; H. = Hatchery.
<sup>3</sup>Reporting Unit = an aggregation of the collections into groups relevant to management and which are accurately resolvable with the genetic data in the baseline. <sup>4</sup>ESU = Evolutionarily Significant Unit; CVFR = Central Valley Fall Run ESU; CVSR = Central Valley Spring Run ESU; CVWR = Central Valley Winter Run ESU; CC = California Coastal ESU; UKTR = Upper Klamath and Trinity Rivers ESU.
<sup>5</sup>N<sub>day</sub>: the number of samples for which the specific sampling day was recorded.

**Figure 1.**
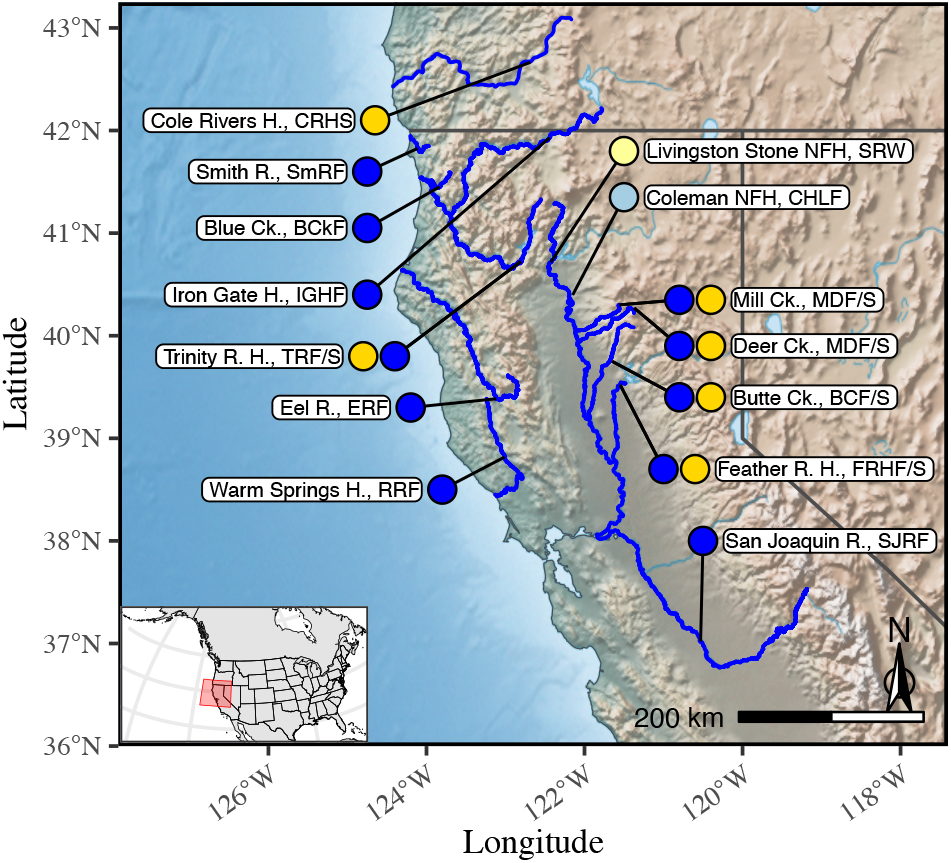
A map of sampling locations represented in the reference baseline. The colored circles aside each location name represent the different run-timing groups of Chinook salmon included in the collections from the location. Color codes are as follows: 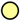= Winter run, 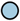= Late-fall run, 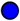= Fall run, 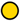= Spring run. Names of each location are followed by the location codes (Table 1). Abbreviations used in location names as follows: R. = River; Ck. = Creek; H. = Hatchery; NFH = National Fish Hatchery.

Collections in each location were separated into fish from the different adult migration timing ecotypes according to a variety of criteria. Winter-run Chinook salmon are propagated at the Livingston Stone National Fish Hatchery for supplementation (released as juveniles into the river) and as a captive broodstock; sampling of this population was performed by hatchery staff during spawning. In Central Valley rivers with both spring-run and fall-run ecotypes (i.e. Mill, Deer and Butte Creeks), previous research has documented different (but overlapping) ranges of spawning time (Julian date) for each ecotype (Fry, 1961; Yoshiyama *et al*., 1998). As such, sampling targeted carcasses found in each of these rivers at times representative of their run-timing designation, which we were able to further confirm using their genotypes within the region of strongest association (RoSA) on chromosome 28 (Thompson *et al*., 2020). In the San Joaquin River, in 2016, at the time of sampling, only fall-run Chinook salmon adults were found, so the samples there were easily categorized. Run-timing designation at the Feather River Hatchery is more complicated but it is managed by the hatchery practices in place: early returning fish in May and April (expressing the spring-run life history) are visibly tagged and returned to the river. Subsequently in the fall, when hatchery raceways are reopened, only fish with the visible tag are included as spring-run broodstock, and the rest are included in the fall-run broodstock. We sampled putative spring-run and fall-run fish from these two broodstocks. Similar to broodstock collections in the CCV, samples from spring- and fall-run fish in the Trinity River were selected on the basis of spawn timing at the Trinity River Hatchery with migration-timing ecotype then confirmed by RoSA genotype. Elsewhere, adult samples were taken from live broodstock at Rowdy Creek (Smith River), Iron Gate (Klamath River) and Warm Springs (Russian River) Hatcheries and juveniles were sampled from Blue Creek (Klamath River); all locations where only fall-run fish have been documented. Spring run adult fish were sampled from the spring-run broodstock at Cole Rivers Hatchery on the Rogue River. Samples from the Eel River were taken from live fish ascending a ladder in the upper basin and were confirmed as fall-run ecotype by RoSA genotype.

In total, the samples in the baseline represent fish from five (Table 1) different Evolutionarily Significant Units (ESUs) of Chinook salmon (Waples,1991).

### Genetic Markers

The genetic markers in the baseline include newly discovered microhaplotypes, SNP assays that were translated into amplicons, markers associated with phenotypes, as well as species- and sex-specific markers. These genetic markers were compiled from several different efforts, as detailed in the sections below.

#### Novel microhaplotype discovery

We identified candidate loci with multiple singlenucleotide polymorphisms (SNPs) among multiple populations using reduced-representation genome sequencing of 2–4 Chinook salmon, each, from a number of populations ranging from California to British Columbia and the interior Columbia River (Table S1). Genomic DNA was extracted using Qiagen DNeasy 96 Blood and Tissue Kits on a BioRobot 3000 (Qiagen, Inc). We used double-digest restriction-site associated DNA sequencing (ddRAD) with different size selections (400 bp and 500 bp fragments) to produce a broad array of locus targets. Library preparation and sequencing methods followed those of Peterson *et al*. (2012) with the modifications described in (Baetscher *et al*., 2018). Sequencing was performed on a MiSeq (Illumina, Inc) using 2 × 300-cycle paired-end sequencing.

Stacks v1.45 (Catchen *et al*., 2013) was used to analyze the ddRAD sequence data. As there was not a well-assembled Chinook salmon genome at the time, a *de novo* assembly was performed. Initial filtering retained RAD loci with two or more SNPs present in at least 10 samples. After this initial filtering, there were 3,931 RAD loci retained in the ddRAD data with 400 bp size selection and 4,794 retained in the data with 500 bp size selection. Gene regions were then selected that (1) contained multiple SNPs within a 100 bp window, and (2) showed haplotype variation amongst the winter run and at least one other population. A 100 bp window was used because the success of multiplex PCR and MiSeq sequencing is diminished in amplicons larger than 100 bp. Primer3 software (Untergasser *et al*., 2012) was used to design primers for 229 gene regions that met these criteria.

These gene regions were PCR-amplified and sequenced on a MiSeq sequencer using a 2 × 75-cycle kit. The sequences from each locus were mapped to the de novo reference sequences (produced by Stacks) with the Burrows-Wheeler aligner using the mem algorithm (Li & Durbin, 2009). Subsequently, alignmets were sorted with SAMtools (Li *et al*., 2009) and provided to FreeBayes (Garrison & Marth, 2012) for variant calling. FreeBayes output was filtered to remove indels and retain variants with a Phred-scaled quality score of 30 or greater and a read depth of 10 or higher. Further filtering and analysis of the loci, including assessment of Hardy-Weinberg equilibrium (Hardy, 1908) was conducted in MICROHAPLOT (Ng *et al*., DOI: 10.5281/zenodo.820110). Loci that produced more than two haplotypes within an individual, as well as those clearly violating Hardy-Weinberg equilibrium within populations were removed.

#### Conversion of existing SNPtype assays

A number of SNP markers from an earlier discovery effort (Clemento *et al*., 2011) had already been validated as useful for GSI and PBT in California (Clemento *et al*., 2014). These markers were originally assayed using TaqMan (Life Technologies Corp., Carlsbad, CA) probes or SNPtype assays (Standard BioTools Inc., South San Francisco, CA). In order to type these markers using amplicon sequencing, we followed the methods in Campbell *et al*. (2015) and added the Illumina small RNA sequencing primer and read 2 sequencing primer to the SNPtype Specific Target Amplification Primer and Locus Specific Primers respectively. We then dropped any loci that produced genotypes that were frequently inconsistent with genotypes produced with the SNPtype assays and any loci that did not amplify well or those that amplified so well that they were grossly over-represented among the reads on the sequencer. We used FreeBayes (Garrison & Marth, 2012) to identify all variants on the fragment (not just the SNPs targeted by the original TaqMan assays). This allows us to score each locus as a microhaplotype (Baetscher *et al*., 2018) when multiple variants occur in the sequence. We filtered out variants that were considered multinucleotide or complex variants by FreeBayes. The resulting genetic markers are hereafter referred to as ‘SNPlicons’.

### Identifying the Late-Fall Associated Region (LFAR) and designing markers therein

To identify loci with large allele frequency differences between late-fall and fall-run Chinook salmon, we first conducted a simple case-control GWAS using the low-coverage whole-genome sequencing data of Thompson *et al*. (2020), mapped to the Otsh_v1.0 Chinook salmon assembly (GenBank accession GCA_002872995.1). We conducted two separate association-study comparisons. The first involved the 16 late-fall-run fish as cases with 16 fall-run fish from the Feather River Hatchery as controls. The second involved the same 16 late-fall fish as cases, but used 16 fall-run fish from the San Joaquin River as controls. The association studies were performed for each chromosome with ANGSD (Kim *et al*., 2011; Korneliussen *et al*., 2014), using the following command: `angsd -yBin` {`ybin`}`-r` {`chrom`} `-minMapQ 30 -minQ 20 -minInd 12 -doAsso 1 -GL 1 -doMajorMinor 1 -doMaf 1 -SNP_pval 1e-6 -out` {`prefix`} `-bam` {`bamlist`}, where {`ybin`} is the path to a file indicating the cases and controls, {`chrom`} indicates the chromosome name, {`prefix`} is the prefix to use for output files, and{`bamlist`} is the path to the text file holding paths to the binary alignment map (BAM) files of aligned sequence data, one for each individual, ordered as in {`ybin`}.

There was only one peak in association scores that was shared between both comparisons (see Results). From this peak we identified candidates for follow-up genotyping by including the 10 SNPs with the lowest association *p*-values, and supplementing those with additional SNPs that showed:

- An estimated absolute allele frequency difference between late-fall and all 32 fall-run fish, |*d*| > 0.75, with an approximate lower confidence interval of |*d*| greater than 0.5, or
- |*d*| > 0.5 with a lower confidence interval >0.25 and with either fall-run or late-fall run being nearly fixed for one of the alleles, or
- An annotation from snpEff (Cingolani *et al*., 2012)of “High” or “Moderate,” or
- |*d*| > 0.48 and a lower confidence interval of >0.25 and a genomic coordinate > 2.2 Mb.

The final condition was implemented to gather several SNPs within the flanking region to the right of the main peak of association.

For the candidate markers found by the above process, we designed amplification primers using Primer3 (Koressaar & Remm, 2007; Untergasser *et al*., 2012) and tested amplification in a subset of Chinook salmon samples. The primer pairs that amplified sequence that reliably mapped back to the association-peak region were then typed in a separate collection of 1,638 Sacramento River Chinook salmon sampled from the fish trap below Keswick Dam, including 1,169 winter run, 281 spring run, and 188 fall/late-fall run. We divided the fish identified from the 125-locus genetic stock identification data set of Thompson *et al*. (2020) as fall/late-fall into fall run and late-fall run according to their sampling date at Keswick: 99 fish arriving before Dec 26 were classified as fall run and 89 fish arriving after Dec 26 were classified as late-fall run. The allele frequencies of the target SNPs at the remaining amplicons were estimated for each of these four groups: winter, spring, fall, and late-fall, and we retained amplicons showing large allele frequency differences between late-fall fish and all other runs.

#### Winter-run associated polymorphisms (WRAPs)

Using the whole genome sequencing data of Thompson *et al*. (2020), we also sought genomic regions that might provide diagnostic markers for the Sacramento winter-run Chinook salmon. This ecotype is highly differentiated from all other Chinook salmon populations throughout its genome, which complicates the process of finding diverged genome regions using a simple association study, with only 16 samples, as done to identify the LFAR. Consequently we pursued a bespoke analysis to find regions we could expect to have large allele frequency differences between the winter-run and all other ecotypes of Chinook salmon in California’s Central Valley. This analysis screened for regions with large allele frequency differences across large blocks of the genome, and is described in the supplemental text section S1.

#### Adult-migration-timing associated markers

We include six amplicons that capture the eight SNPs within the “Region of Strongest Assocation” (RoSA) between spring-run and fall-run fish, near the ROCK1 and GREB1L genes, on Chromosome 28 listed in Table S3 of Thompson *et al*. (2020).We add three more amplicons in the same RoSA region to the panel, as well as one more amplicon that allows genotyping of snp670329, which was discovered and described by Thompson *et al* (2019). Including all these markers in the panel allows them to be routinely typed, creating a database for understanding the spatial and temporal distribution of the alleles in this genomic region; however we have omitted these markers for calculating likelihoods for GSI, since their distribution across populations in the Feather River has been clearly and evidently influenced by hatchery practices (see Results), it is not necessary to include them for accurate assignment, and because the 12 SNPs within these three amplicons are typically in profound linkage disequilibrium (LD) making them inappropriate for use as separate markers in most software for population assignment or GSI, that typically assumes markers are not in LD within the component populations of the baseline.

#### Additional phenotypically associated markers

The baseline also includes three amplicons with genetic variation in two genes, VGLL3 and SIX6, that have been found to be associated with phenotypes related to reproductive maturity in both Atlantic (Barson *et al*. 2015) and Pacific (Waters *et al*., 2021) salmonids. We include them both to provide additional variation for population genetic analyses and to monitor populations for potential phenotypic patterns.

We include an assay from a novel locus that targets the sex-determining region (*sdY*) on the Y chromosome in Chinook salmon. We designed this assay to target the SNP described by Bertho *et al*.(2022)that differentiates the functional *sdY* and a nonfunctional *sdY* pseudogene. Examination of whole-genome sequence data in multiple populations of California Chinook salmon (Thompson *et al*., 2020) shows that this marker characterizes genetic sex in diverse California salmon populations more accurately than previously published assays (Von Bargen *et al*., 2015). And, finally, we include marker Ots_coho001_05_32691399 which is fixed for alternate alleles in coho salmon and Chinook salmon. This is helpful when samplers have unwittingly collected coho salmon.

### Localizing markers within the genome

The novel microhaplotype markers and the SNPlicon targets were developed prior to the advent of a well-assembled genome for Chinook salmon. As a consequence, many of these markers had been used previously without certainty about their location in the genome. After designing and validating an amplicon-sequencing approach to type both of these groups of markers we identified their locations within the genome by mapping the consensus sequences (roughly 100–300 bp) around the markers to both the Chinook salmon version 1 (Otsh_v1.0, GenBank accession GCA_002872995.1) and version 2 (Otsh_v2.0, GenBank accession GCA_018296145.1) genome assemblies, using both bwa-mem (Li & Durbin, 2009) and the BLAST-like Alignment tool (Kent, 2002). A similar mapping exercise was performed for the remaining markers so that we can report their locations in both Otsh_v1.0 and Otsh_v2.0.

### Amplicon sequencing, microhaplotype calling, and further quality control

Amplicon sequencing was performed with the Genotyping-in-Thousands-by-sequencing (GT-seq) method (Campbell *et al*., 2015). All loci were multiplexed with primer concentrations ranging from 0.083 uM to 0.5 uM to increase uniformity of read depths across all loci. Sample normalization was performed using the SequalPrep Normalization Plate Kit (Applied Biosystems) according to the manufacturer’s instructions. Samples were pooled by plate and then purified using Agencourt AMPure XP magnetic beads. Libraries were quantified either by qPCR with the NEBNext Library Quant Kit for Illumina (New England BioLabs) or with the Qubit dsDNA HS Assay Kit (Thermo Fisher Scientific) before dilution and pooling for sequencing. All samples were sequenced on a MiSeq using 2 × 75-cycle paired-end sequencing protocols. Primer sequences and concentrations for making the primer pool are in Supplemental Data 1.

For evaluation of candidate loci, dual-barcoded sequences were used to identify individual tissue samples using the MiSeq analysis software (Illumina). The paired-end sequence reads were joined together using Fast Length Adjustment of SHort reads (FLASH: Magoč & Salzberg, 2011)) and mapped to an indexed sequence reference created from the Stacks target loci (for the novel microhaplotypes, SNPlicons and RoSA markers) or from the entire Otsh_v1.0 genome (for the LFAR, WRAP, VGLL3, and Six6 markers) using BWA mem(Li & Durbin, 2009). Mapped reads were converted to BAM files using SAMtools (Li *et al*., 2009) and FreeBayes (Garrison & Marth, 2012) was used to call variants.

For genotyping of baseline samples, final filtering at each locus required a read depth of 20 and a read depth ratio of 0.30 (i.e., for heterozygotes the lower read depth haplotype must have 30% of the total reads at that locus). MICROHAPLOT was used to call alleles and genotype processing was performed with R scripts.

Each locus was assessed for the presence of null alleles using the R package ‘whoa’ (https://github.com/eriqande/whoa; seeHendricks *et al*., 2018).

### Population genetic analyses

Basic data summaries were calculated for each population/collection in the reference baseline using simple operations in the R programming language, Version 4.3 (R Core Team, 2024). We started with a simple tally of the number of alleles found at each locus. To provide a summary of the genetic variation and missing data we further summarized:

- the average over loci of the number of fish in which each locus was successfully genotyped,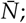
- the average number of alleles per locus in each collection with all collections downsampled to the smallest number of genotyped individuals at each locus,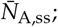
- the fraction of polymorphic loci in each collection after subsampling to the smallest number of genotyped fish,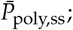
- the expected heterozygosity as the average over loci of one minus the frequencies of homozygotes at each locus expected from the estimated allele frequencies,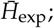 and
- the observed heterozygosity as the average over loci of the fraction of heterozygous genotypes,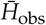.

When subsampling each collection to the minimum number of genotyped fish across all collections at a locus, we subsampled individuals without replacement, taking the average of 1,000 different random subsamples. We calculated Weir & Cockerham (1984)’s pairwise *F*_ST_ between all collections, using the `pp.fst` function from the R package ‘hierfstat’ (Goudet & Jombart, 2022). Finally, we evaluated population structure in the data by performing unsupervised clustering using the program STRUCTURE (Pritchard *et al*., 2000; Falush *et al*., 2003). For this analysis, we omitted the run-timing associated markers. We performed 20 replicate runs at each value of the assumed number of subpopulations, *K*, in {2, 3, 4, 5, 6, 7} using default settings of the program. STRUCTURE output was a summarized and visualized using CLUMPAK (Kopelman *et al*., 2015).

### Power for genetic stock identification and population assignment

To assess the power of the marker panel for GSI, we used the R package ‘rubias’ (Moran & Anderson, 2019). We used the function `self_assign()` to perform assignment of each individual in the reference baseline to one of the collections within the baseline. Each fish was removed from the reference baseline when calculating the likelihood that it originated from the collection it actually belongs to, using a classical leave-one-out approach to eliminate the inflated estimates of power that might occur without such a procedure (Anderson *et al*., 2008). The scaled likelihoods of collection membership returned by `self_assign()` were summed within reporting units, and fish were assigned to the reporting unit with the largest value of that sum. Subsequently, fish were assigned to the collection within that reporting unit with the highest scaled likelihood. We considered two thresholds for assignment. In the first, all fish are assigned, in the second only fish with a summed (i.e., summed over all collections in a reporting unit) scaled likelihood greater than 0.8 were assigned.

Finally, we investigated the matrix of assignments and misassignments broken down according to the genotype of the fish at the markers within the RoSA. For this we identified the RoSA genotype (EE, EL, or LL, see Thompson *et al*., 2020) from the concordance of the genotypes at all 8 markers described in (Thompson *et al*., 2020), requiring that no genotypes be missing within an individual from any of those 8 markers, and discarding data from 12 individuals in which the E or L haplotypes had clearly recombined.

### Power for relationship inference

To explore the utility of the marker panel for relationship inference, we used the R package ‘CKMRsim’ (https://github.com/eriqande/CKMRsim) to estimate the false-positive rates (FPRs) expected at different values of the false negative rate (FNR) for a variety of different pairwise relationships in different populations. For these purposes, we divided the collections into 10 different groups. The members of each group are relatively, genetically similar, but the groups do not necessarily correspond to reporting units, because, in some cases, we desired FPRs and FNRs for a single collection. For all cases, we assumed that the genotyping error rate was 1% per locus and the genotyping error model was the default “True-genotype-independent” model.

For errors involving misidentifying unrelated (U) pairs as parent-offspring (PO), full-sibling (FS), or half-sibling (HS) pairs we used importance sampling to estimate the very small FPRs. The FNRs were estimated in each case using simple Monte Carlo. For the FS and HS cases the FNRs were calculated from Monte Carlo sampling while taking account of the physical linkage of the markers by inputting their genomic positions into ‘CKMRsim’ and using the package’s interface to the MENDEL (Lange *et al*., 2013) software to simulate genotypes of related pairs in the presence of physical linkage.

For PO pairs (and U pairs), FNRs were calculated without reference to the genomic locations of the markers because physical linkage does not affect the distribution of log-likelihood ratios in those two relationship types, because there is no variation, anywhere in the genome, in the number of gene copies shared identical by descent between a parent and its child or between two unrelated individuals.

For errors involving misidentifications between the relationship types of avuncular (AN: aunt-niece, uncle-nephew, etc.), PO, FS, and HS, we used regular Monte Carlo to estimate the FPRs and FNRs, because importance sampling cannot be used in those cases. Since there are expected to be far fewer such relationships than the number of unrelated pairs, the relevant FPRs are large enough that they can be reliably estimated (at least down to rates of 10^−3^) without importance sampling. For AN, FS, and HS, both the FPRs and FNRs were estimated while taking account of physical linkage of the markers, as described above.

## Results

### Genetic Markers

A summary, which includes genomic location, consensus sequence, and primer sequences, of all the amplicons used in the baseline is available in Data Supplement 1. An overview of genomic coordinates is found in Fig. S2

#### Additional microhaplotype discovery

We tested the 229 candidates from our novel microhaplotype discovery process and initially retained the 125 markers described in (Thompson *et al*., 2020) for use in stock identification within the Klamath Basin. However, further evaluation of those markers in other populations led us to remove some of the markers. Twelve loci appeared to have null alleles in the winter-run population and were removed. Additionally, two loci were monomorphic in the winter-run, even though they contained multiple alleles in other populations, and an additional 5 amplicons were deemed difficult to score in routine application of the baseline, and were removed, leaving 106 novel microhaplotype amplicons in the current panel.

#### Conversion of existing SNPtype assays

Of the 96 existing SNPtype assays, we were able to convert 78 into reliable amplicon-sequenced assays that were included in the panel.

#### Late-fall Associated Region Markers

The case-control association test revealed a single peak on chromosome 34 with large allele frequency differences between Central Valley late-fall and fall-run fish.

This peak was evident both in the comparison of late-fall fish to fall-run fish in the Feather River Hatchery and of late-fall fish to the fall-run fish in the San Joaquin River (Figure S1).

The Manhattan plot of Figure S1 shows sporadic SNPs with low association *p*-values. These are not shared between the two different late-fall to fall comparisons, and sites that we investigated were concentrated in areas of poor mapping, suggesting that their *p*-values were artifactual. By contrast, a large number of sites near the prominent peak on chromosome 34 had large allele frequency differences between late-fall and fall run. Taking the 10 SNPs with the lowest association *p*-values, and including additional ones from the filtering criteria given in the Methods, yielded 49 candidate markers for which we designed primers for amplication. Only 8 of these 49 primer pairs yielded reliable amplification and mapping. Of these 8, two of the SNPs showed inconsistent genotyping and were removed from consideration. Of the remaining 6, only 3 showed marked allele frequency differences (> 0.7) between late-fall and fall run. These three SNPs (located at: Chr34:828,768, Chr34:865,057, and Chr34:1,063,084 in the Otsh_v1.0 assembly) also had pronounced differences in allele frequency between late-fall and spring or winter run.

Analysis of the mapping of amplicons for Chr34:828,768 showed that many (≈89%) of the reads were off-target (aligning to other chromosomes, etc.), making it costly (in terms of sequencing effort) to include the marker in the baseline panel. Consequently it was dropped from the panel. The two remaining loci, which are at coordinates Chr34:865,057, and Chr34:1,063,084 in the Otsh_v1.0 assembly are at coordinates Chr34:954,054 and Chr34:1,151,868 in the Otsh_v2.0 assembly, and the frequency of the different alleles at the two remaining loci within different reporting units in the baseline are shown in Table 2.

**Table 2.** Allele frequencies across reporting units of the two late-fall associated markers. Frequencies are given for the alleles most common amongst late-fall Chinook: nucleotide `G` at locus Chr34:954,054 and `A` at Chr34:1,151,868. *N* is the total number of gene copies available for estimating the allele frequency. Genome coordinates of SNPs are from Otsh_v2.0.

| Reporting Unit | Chr34:954,054 |  | Chr34:1,151,868 |  |
| --- | --- | --- | --- | --- |
|  | freq. | *N* | freq. | *N* |
| CV-Late-Fall | 0.750 | 516 | 0.733 | 592 |
| CV-Fall | 0.054 | 910 | 0.030 | 908 |
| CV-Spring | 0.023 | 396 | 0.003 | 396 |
| CV-Winter | 0.005 | 222 | 0.005 | 222 |
| Cent-Cal-Coast | 0.000 | 92 | 0.004 | 278 |
| Klamath-Trinity | 0.000 | 172 | 0.000 | 358 |
| SO-NCal-Coast | 0.000 | 192 | 0.000 | 304 |

#### Winter-run-associated polymorphisms

Though the winter run fish are easily identified, we undertook an additional effort to try to find markers that could be diagnostic for the winter run, distinguishing the winter run from all other groups of Chinook salmon in the CCV. Initial examination of sequence data in the 16 winter-run fish and 84 non-winter run fish revealed a number of SNPs that were potentially diagnostic, but none were fixed for alternative alleles in a larger sample. Nonetheless, three of these markers from that effort had large frequency differences between the groups and are included in the reference baseline baseline. The process of discovering them is described in the supplemental text section S1.

### Localizing markers within the genome

Of the 184 loci that originated from our novel microhaplotype discovery or from SNPtype assays, 179 of them mapped identically to a single location on an assembled chromosome in the Otsh_v2.0 genome, using both bwa mem and BLAT. Of the remaining 5 loci, one of them was mapped to the same location by BLAT and bwa mem, but the alignments differed in length, two of them had a single secondary alignment on a different chromosome that was identical in both bwa mem and BLAT, one of them had only a fragment mapping to the secondary alignment, and one of them had multiple non-primary alignments (Supp. Data 1).

Mapping to Otsh_v1.0 was similar, except that a greater number of markers mapped to unplaced scaffolds in Otsh_v1.0 than in Otsh_v2.0, likely reflecting the more complete nature of the second assembly.

### Population genetic summaries

The total number of alleles per locus varied between 1 and 10 (Figure S3), with the only locus bearing a single allele in Chinook salmon being the species-specific marker Ots_coho001_05_32691399. Biallelic loci were most common, at 66 of the total, but two-thirds of the microhaplotype markers showed more than two alleles, which has been shown to provide significantly more power for relationship inference than a comparably sized panel of biallelic SNPs (Baetscher *et al*., 2018).

The population-genetic summary statistics are presented in Table 3. The average number of fish with non-missing data in each population varied from a low of 40.2 (TRF) to a high of 279.6 (CHLF). The average number of alleles per locus, standardized to the smallest sample size, ranged from 2.32 (SRW) to 2.64 (RRF), with SRW also having the smallest value of the standardized fraction of polymorphic markers at 0.88 and RRF sharing the highest value (0.96) with FHRF and BCkF. SRW also had the lowest values for expected and observed heterozygosity: 0.338 and 0.329, respectively. While the highest for expected heterozygosity was 0.401 (SmRF) and for observed heterozygosity, 0.399 (CRHS).

**Table 3.** Simple population genetic summaries. All quantities are means over loci. 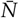 is the number of fish with observed genotypes; 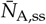 is the average number of alleles and 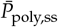 the fraction of polymorphic loci, after sub-sampling to the smallest sample size per locus; 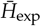 and 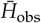 are expected and observed heterozygosity, respectively. Population codes are as given in Table 1

| Code | $\bar{N}$ | $\bar{N}_{A,ss}$ | $\bar{p}_{poly,ss}$ | $\bar{H}_{exp}$ | $\bar{H}_{obs}$ |
| --- | --- | --- | --- | --- | --- |
| CRHS | 49.6 | 2.60 | 0.95 | 0.398 | 0.399 |
| SmRF | 45.0 | 2.60 | 0.95 | 0.401 | 0.394 |
| BCKF | 54.0 | 2.59 | 0.96 | 0.393 | 0.382 |
| IGHF | 90.9 | 2.37 | 0.93 | 0.356 | 0.363 |
| TRS | 44.8 | 2.37 | 0.92 | 0.353 | 0.348 |
| TRF | 40.2 | 2.37 | 0.92 | 0.352 | 0.351 |
| ERF | 91.5 | 2.51 | 0.95 | 0.364 | 0.356 |
| RRF | 89.5 | 2.64 | 0.96 | 0.398 | 0.391 |
| SRW | 73.4 | 2.32 | 0.88 | 0.338 | 0.329 |
| BCS | 110.6 | 2.52 | 0.92 | 0.379 | 0.373 |
| MDS | 77.1 | 2.55 | 0.94 | 0.394 | 0.397 |
| FRHS | 180.1 | 2.61 | 0.95 | 0.397 | 0.392 |
| FRHF | 103.9 | 2.63 | 0.96 | 0.394 | 0.398 |
| BCF | 40.3 | 2.60 | 0.94 | 0.385 | 0.370 |
| MDF | 64.4 | 2.59 | 0.94 | 0.386 | 0.383 |
| SJRF | 78.0 | 2.57 | 0.94 | 0.384 | 0.380 |
| CHLF | 279.6 | 2.53 | 0.94 | 0.386 | 0.384 |

The estimated pairwise *F*_ST_ values between the collections varied from a low of 0.0 to a high of 0.288 (Figure 2a). Notably, most of the largest values of *F*_ST_ occurred between SRW and another collection.

**Figure 2.**
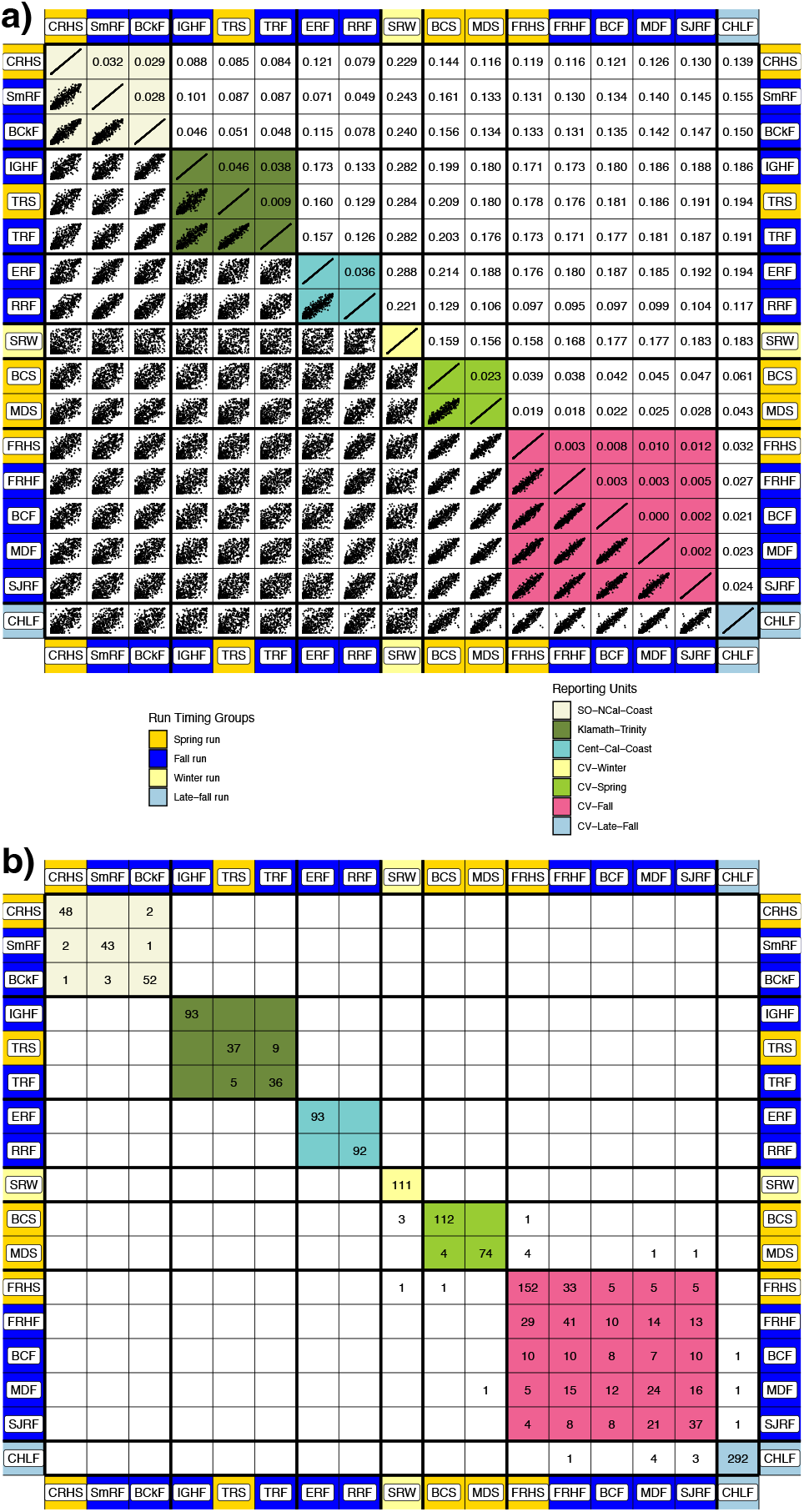
(**a)** Pairwise *F*_ST_ values and allele frequencies, and (**b)** self-assignment rates. The codes of the populations appear on the outer margins of the table atop cells colored according to the run-timing group of the collection. Interior cells are colored according to reporting unit. In (**a**), for *F*_ST_ values, the upper triangle was computed using all the genetic markers except for those in the RoSA on chromosome 28. Each cell in the lower triangle shows a small scatter plot of estimated allele frequencies in the collections—the *x* value is allele frequencies from the collection in the column and the *y* value is from the collection in the row. Frequencies for all alleles are shown, with values within each cell ranging from 0 to 1. These frequencies were estimated by, for each locus, randomly subsampling individuals from each collection to the minimum number of observed alleles in any collection at the locus. In (**b**), for self-assignment tallies, the row corresponds to the true source collection and the column corresponds to the assigned collection. All individuals were assigned here, using a maximum-scaled-likelihood rule to reporting unit and then to collection within reporting unit as described in the text. For the assignment matrix resulting when only fish with a scaled likelihood > 0.80 to reporting unit are assigned, see Figure S5

At values of *K* from 2 to 7, clusters identified by the program STRUCTURE generally corresponded to groupings of related populations, and confirmed *a priori* knowledge about which collections in the baseline could and should be grouped together into reporting units (Figure 3). At every value of *K*, CLUMPAK discerned at least 12 of 20 replicates in the major mode (Figure 3). Clustering solutions in the minor modes generally converged on a single alternative solution and appear in Figure S4. The late-fall run collection (CHLF) appears as a separate cluster at *K* values as low as 4, with the inclusion of the LFAR markers. This is notable, since accurately discriminating between the late-fall and fall-run Chinook salmon of the Central Valley has been impossible with previously used genetic marker sets. Another noteworthy feature appears at *K* = 7, with STRUCTURE separating the fall-run reporting unit into two clusters (orange and red) that do not correspond exactly with the individual collections within the reference baseline. For example, fish from the orange cluster that predominates in the FRHS collection are also found in the FRHF and MDS collections.

**Figure 3.**
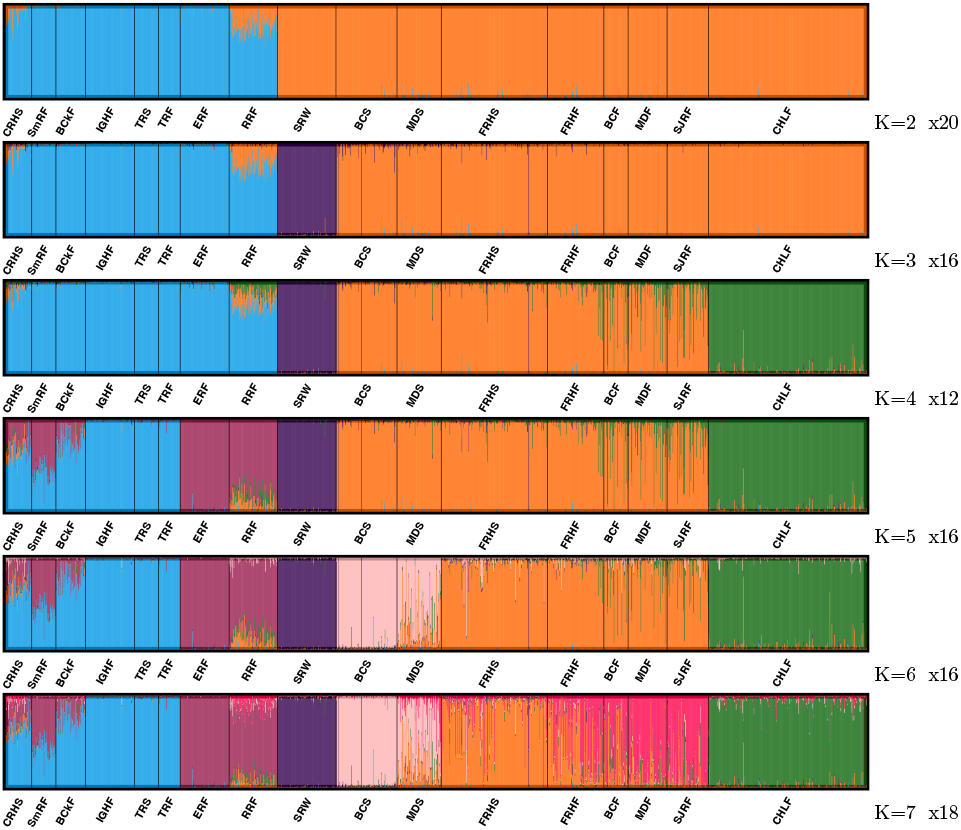
Major modes found by STRUCTURE and summarized by CLUMPAK out of 20 replicates for each value of *K*, the assumed number of subpopulations. *K* value is listed at lower right of each barplot, followed by the number of replicate runs of structure that converged to this major mode.

### Power for genetic stock identification and population assignment

Leave-one-out cross-validation (self-assignment) analysis demonstrates that this reference baseline provides a high degree of accuracy for distinguishing the major groups of Chinook salmon in California. The results of the analysis are summarized in a self-assignment matrix (Figure 2b). This figure shows the collections divided into the seven different reporting groups: SO-NCal-Coast, Klamath-Trinity, Cent-Cal-Coast, CV-Winter, CV-Spring, CV-Fall, and CV-Late-Fall. These groupings correspond largely to the divisions in the data found by STRUCTURE, and they also correspond with divisions amongst the stocks as defined for management, and, as seen in Figure 2b, they correspond with groups of populations that can be reliably distinguished from one another for population assignment.

Overall, of the 1636 fish in the reference baseline, 1612 (98.5%) were correctly assigned to their reporting unit of origin, while 24 (1.5%) were incorrectly assigned. Four of those 24 misassignments which involved fish being incorrectly assigned to SRW, were clearly fish from SRW that had been incorrectly sampled into a different collection, likely due to straying. (This is evident because SRW are so distinct that it is highly improbable they would be incorrectly identified). Of the remaining 20 misassignments, 8 involved fish from the CHLF (late fall run) collection being misassigned to the CV-Fall reporting unit and 6 were fish from the MDS (Mill-Deer spring run) collection being assigned to the CV-Fall reporting unit. It is possible that some of these misassignments represent CV-Fall fish that were incorrectly sampled as CHLF or MDS; however, unlike with SRW, we cannot conclude that confidently, as there is some degree of overlap in likelihood values between the two.

While fish from CHLF were found to misassign at a rate of 2.7% (8/300) and MDS at a rate of 7.1% (6/84) to CV-Fall, the converse is not evident—that is, fish from the CV-Fall are misassigned to CHLF at a rate of only 0.6% (3/508) and to the CV-Spring reporting unit (to which MDS belongs) at a rate of only 0.4% (2/508).

When assignments are only accepted with a scaled likelihood greater than 0.8, 20 assignments are discarded, and half of those (10) were the incorrect assignments. Thus, at the 0.8 threshold, the total fraction of correct assignments is 99.1% (Figure S5). Notably with the >0.8 criterion, the misassignment rates from the abundant CV-Fall reporting unit to CV-Late-Fall or to CV-Spring both drop below 0.2% (1/502).

As seen in previous studies, FRHS, the Feather River Hatchery spring-run stock, is considerably differentiated from the remaining spring-run stocks in the Central Valley (MDS and BCS). This is reflected in the low misassignment rates of FRHS fish to the CV-Spring, and is also evident in the STRUCTURE results. We note, however, that the RoSA markers provide the differential signal necessary to distinguish Feather River spring-migrating fish from the other CCV stocks with a predominantly fall-run genomic background.

FRHS fish are produced by selecting spring-tagged fish that again return in fall, when the hatchery ladder reopens. This results in untagged spring run, fish not encountered during the spring trapping season, being used as broodstock for the fall-run program. This is evident when the assignment matrix is enumerated in terms of RoSA genotypes (Figure S6). In this context, it is clear that the “fall-run” hatchery program in the Feather River generates many spring-run fish and RoSA heterozygotes. Similarly, in other regions, such as the Trinity River, the RoSA markers are the only reliable means to distinguish fall- and spring-run fish, as they otherwise share genomic background.

In California, only the Sacramento and Klamath basins have documented spring-run salmon and their historical occurrence in other basins has been unclear; however we identified three additional copies of a recombinant haplotype that carries half of the RoSA SNPs from both the E and L lineages in the Eel and Russian rivers. No other clear recombinants were identified in the study populations, suggesting that it arose in the California Coastal Chinook Salmon lineage, and providing further evidence of past presence of RoSA haplotypes associated with early migration.

### Power for relationship inference

The ‘CKMRsim’ power analyses for this set of markers demonstrates that they have ample variation for accurate identification of parent-offspring (PO) and full-sibling (FS) pairs in almost all realistic situations. The predicted false positive rates from unrelated pairs for both the parent-offspring (PO) and full-sibling (FS) relationships are exceedingly low for all population groups, even when using stringent false negative thresholds as low as 0.05 (Figure 4a). For example, the false positive rate for FS identification is less than one error in 10 million comparisons of potential siblings in the California Chinook salmon population with the lowest heterozygosity (SRW) at a false negative rate of 0.05.

**Figure 4.**
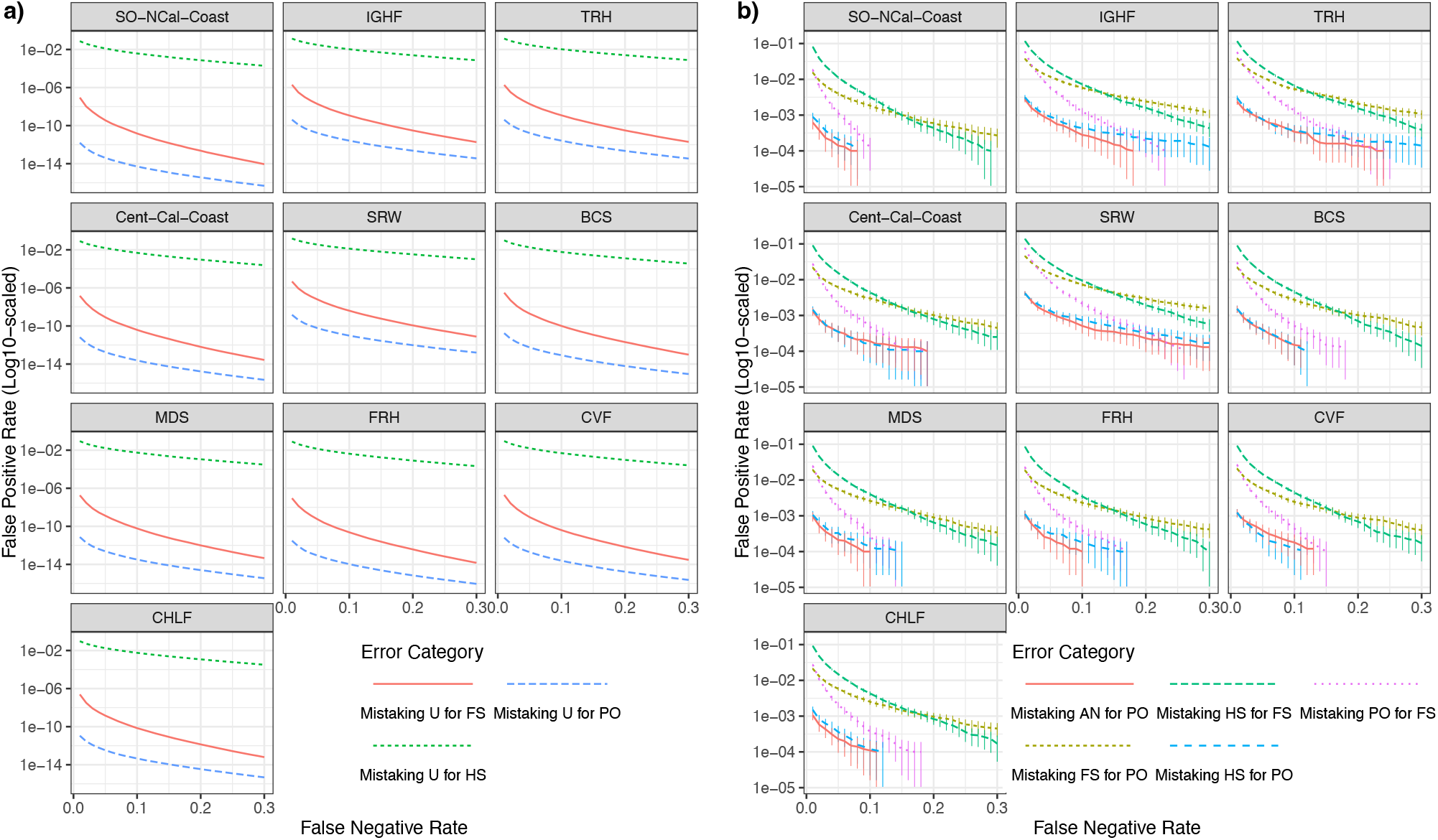
False positive rates (FPRs) as a function of false negative rates (FNRs) for distinguishing between different relationships. Groupings for panels that are neither a previously defined reporting unit nor a single collection (Table 1) are as follows: TRH = {TRS, TRF}; FRH = {FRHS, FRHF}; CVF = {BCF, MDF, SJRF}. **(a)** FPRs calculated using importance sampling of unrelated pairs of individuals being incorrectly categorized as parent-offspring (PO), full-sibling (FS) or half-sibling (HS) pairs. **(b)** FPRs calculated using regular Monte Carlo sampling of avuncular pairs (AN: aunt-niece, uncle-nephew, etc.) or full-siblings (FS) or half-siblings (HS) being incorrectly categorized as parent-offspring (PO); as well as FPRs of parent-offspring (PO) pairs or half-sibling (HS) pairs being incorrectly inferred as full siblings (FS). In both *a* and *b*, 100,000 Monte Carlo replicates were simulated. Importance sampling allows estimation of very small false positive rates for unrelated pairs, but cannot be applied to truly related pairs. Consequently, it is difficult to accurately estimate small probabilities (10^−4^ or less) in **b**, as indicated by the error bars, which represent the estimate ± 2*s* where *s* is the estimated standard error of the mean of the Monte Carlo sample. This also is the reason that many of the lines terminate.

Half-sibling (HS) pairs cannot be accurately distinguished from unrelated pairs in any populations without very large false negative rates and in very modest-sized studies with very few pairwise comparisons.

In a reasonably large population, the vast majority of pairs of individuals are expected to be unrelated, or at least, effectively unrelated, sharing common ancestors only many generations in the past. However, a small fraction of pairs will be more recently related, and distinguishing these kin pairs from target relationships (like PO and FS) must be accounted for. ‘CKMRsim’ provides facilities for assessing error rates between different kin groups, and results for these are summarised in (Figure 4b). Here, again, focusing on the results for the genetically depauperate SRW, and using an FNR of 0.1 we see that the chance of incorrectly categorizing an HS as an FS, or an FS as a PO is less than 1 in 100. Likewise, we can expect PO to be misidentified as FS at a rate of around 1 in 500, and HS or Aunts/Uncles to be misidentified as parents at a rate less than 1 in 1000. Though these rates are much higher than the FPRs for unrelated pairs, they should be compared to the number of actual non-target kin pairs expected. For example, in a salmon population, the number of full aunts or uncles of a fish is unlikely to exceed 50, at later life stages, so a FPR of 0.001 is comfortably low in that case.

The presence of multiple alleles at many of these loci improves power for kin inference compared to a marker panel using only a single SNP at each amplicon. To provide a visual display of the difference afforded by using microhaplotypes compared to single SNPs in this data set, we show the distribution of log-likelihood ratios for different relationships using our microhaplotype-scored amplicons, versus using the most heterozygous single SNP from each amplicon (S7). The figures show a considerable increase in separation between the relationships categories from calling genotypes as microhaplotypes in these amplicons.

While there is not a currently available method (like that of Anderson & Garza, 2006)) for estimating error rates in inference of parent-offspring trios with multiallelic marker data, given the substantial amount of statistical power for identifying PO and FS pairs, which require much more statistical power, it is clear that PO-trio inference with these markers will be essentially error free.

## Discussion

We describe a novel panel of microhaplotype genetic markers for Chinook salmon in California that provides sufficient power for highly accurate identification of parent and offspring pairs and full siblings, and also allows near-perfect identification of individuals to population or genetic group of origin in California. This set of genetic markers includes multiallelic gene regions with high variability for relationship inference, gene regions identified in whole-genome sequence data for increased power for identification of specific populations, and markers that have long been in use for both GSI and PBT in California (Clemento et al., 2014), by converting them into so-called ‘SNPlicons.’ This set of markers lays the foundation for a comprehensive genetic monitoring and evaluation effort that facilitates multiple types of inference and is flexible and extensible.

This marker set and baseline reference dataset provide unprecedented power for identifying fish from all of the Chinook salmon ESUs in California, as well as individual populations within those ESUs (Figure 2). The California Central Valley (CCV) is one of the largest river basins on the west coast of North America and drains the Sierra Nevada and southern Cascade mountain ranges. It has the highest diversity of recognized Chinook salmon ecotypes in the species range, with four named ecotypes, two of which are protected under the US Endangered Species Act. It is also the source of household water for tens of millions of California residents and millions of acres of arguably the most productive and valuable agriculture area in North America. As such, accurate identification of the distinct ecotypes of CCV Chinook salmon is of utmost importance for monitoring and evaluation of individual ecotypes, as well as designing and implementing effective conservation and management actions. This has been challenging, given the recent common ancestors of these ecotypes (Clemento et al., 2014), and ongoing migration between subbasins where different ecotypes predominate.

We describe the first set of genetic markers that produces easily replicable data and that identifies all of the ecotypes in the CCV, including the late-fall-run Chinook salmon ecotype. The late-fall-run occurs only in the CCV and shares a genomic background with the more common CCV fall-run salmon ecotype, so has been refractory to previous GSI efforts with other marker types (Seeb *et al*., 2007; Clemento *et al*. 2014; Meek *et al*., 2016; Thompson *et al*., 2024).

Moreover, we show how the FRH spring-run lineage is easily identifiable through a combination of traditional GSI and the characterization of functional genetic markers in the RoSA. Finally, although previous work has described GSI capabilities that distinguish the natural-spawning CCV spring-run lineages from each other and their fall-run counterparts with moderately high accuracy, we demonstrate near complete accuracy in distinguishing these ‘stocks’. Moreover, the few fish that are apparently misidentified (Figure 2) are likely to be primarily migrants and not true misidentifications. For example, the three fish that were field-characterized as spring-run from Butte Creek, but are genetically identified as winter-run salmon clearly carry the winter-run genomic background and are strays from the winter-run stock, as these could not realistically be misidentified on the basis of the genetic data.

Results from the model-based clustering analysis with the program STRUCTURE, revealed patterns of population relationships that are coincident with previous work (Clemento *et al*., 2014; Kinziger *et al*., 2013) emphasizing the distinction between Chinook salmon populations in Coastal California and the Central Valley. These analyses also uncover some additional patterns, including the clear presence of mixed ancestry in the Coastal California population in the Russian River, which is consistent with its location, at the southernmost edge of the Coastal Chinook salmon distribution and proximate to the mouth of the Sacramento River (the Golden Gate). In the Central Valley, the clustering results emphasize the genetic distinction of the SWR population, which is likely at least partially due to the extreme bottleneck that it passed through in the 1980s and 1990s (Hedrick & Hedgecock, 1994). The inclusion of the LFAR markers also resulted in the clear distinction of the CHLF group, which is coincident with the long-known phenotypic distinction of this population, but represents a novel genetic result. Moreover, at K = 7, the fall-run reporting group breaks into two clusters, which are distributed across almost all of the spring and fall-run populations in the Central Valley, albeit not equivalently, emphasizing both current and historical migration and gene flow between them.

Examination of the geographic patterns of allele frequencies for the RoSA-associated loci found a clear instance of the early-migrating haplotype present in the California Coastal Chinook salmon lineage. This is of note because this lineage does not currently have an early migrating component, although it has long been speculated that the Eel River, at least, historically harbored early-migrating Chinook salmon, and early migrating steelhead (*O. mykiss*) persist in the basin(Bjorkstedt *et al*., 2005). Moreover, we discovered several recombinant RoSA haplotypes in both of the CCC populations that indicate a long-standing presence of both the early- and late-migrating haplotypes in these populations. This is consistent with the idea that early-migrating populations arise through migration of small numbers of individuals carrying E-lineage haplotypes, in either heterozygous or homozygous form, which are then positively selected when appropriate habitat conditions exist. A similar pattern of recombination, due to long-standing co-occurence of the E and L haplotypes in heterozygotes, has been previously documented in the Klamath River (Thompson *et al*., 2020).

Previous genetic marker sets for California Chinook salmon had sufficient power for inferring parent-offspring relationships, but only when both parents were sampled and genotyped. This marker set considerably increases the capacity for relationship inference in California salmon, by providing sufficient power for parent-offspring relationship inference, when only one parent is sampled, as well as identifying pairs of full siblings when no parents are sampled. Groups of full siblings larger than two can be identified even more accurately than expressed in Figure 4, when using statistical approaches that account for the joint relationships between more than two individuals, such as COLONY (Wang, 2004). Although some additional errors might be expected when related but non-target kin pairs are sampled, the false positive rates for these non-target kin pairs are sufficiently small for almost all realistic scenarios. Accurate relationship inference in even the most genetically depauperate Chinook salmon population in California and rangewide (Seeb *et al*., 2007; Clemento *et al*., 2014) is therefore possible with this marker set. We note that, as in previous work, it is nearly impossible to identify HS pairs accurately with data from this, or any standard marker dataset. Accurate identification of HS pairs typically relies on many hundreds of micro-haplotype markers of SNP markers (Baetscher *et al*., 2018) or thousands (Hillary *et al*., 2018).

As sequencing costs drop, whole genome sequence (WGS) data may become the preferred data type for salmon genetics. If so, it may become possible to include all the >200 genetically distinct Chinook salmon populations in North America within a single standardized reference baseline constructed with WGS data that performs equally well at broad and regional scales (DeSaix *et al*., 2024). Such an approach would be highly flexible and extensible, as it would allow for the assignment of unknown-origin fish using just about any marker type, including reduced representation DNA sequencing (e.g., RADseq, Meek *et al*., 2020; Thompson *et al*., 2024), as the variation used by such region-specific panels of markers should be contained within the WGS baseline dataset. For the present, however, baselines tailored to specific regions are essential for regional management questions and targeted sequencing approaches have proven to be the most practical for large-scale applications of either GSI, PBT or an integrated monitoring program (Beacham *et al*., 2021).

## Acknowledgements

Many people from various agencies and watershed groups contributed to the collection of the samples analyzed here, including staff from the California Department of Fish and Wildlife, National Marine Fisheries Service, US Fish and Wildlife Service, US Army Corps of Engineers and Pacific States Marine Fisheries Commission. We also thank Elena Correa, Nicole Anderson and Libby Gilbert-Horvath for contributions to data collection. The manuscript was improved by comments from Jeff Rodzen. USDA is an equal opportunity provider and employer. Mention of trade names or commercial products in this publication is solely for the purpose of providing specific information and does not imply recommendation or endorsements by the U.S. Department of Agriculture.

## Data Accessibility

All data and code needed to reproduce the results here are available online.

- Online version of data and scripts used in paper: https://github.com/eriqande/california-chinook-microhaps
- Archived version of data and scripts used in paper: zenodolinkwhenmade.
- Data repository with full baseline data set. UPDATED WHEN JOURNAL PROVIDES DOI. https://dryad.something.or.other

## SUPPLEMENT TO

### S1 Winter-run-associated polymorphism methods and results

We used the whole genome sequencing data from (Thompson *et al*., 2020) to seek variants with large allele frequency differences between winter-run Chinook salmon and all the other Chinook salmon ecotypes in the Central Valley of California (CCV). Because winter-run Chinook salmon are already highly differentiated from all others, we did not pursue an association study as used for identifying the late-fall-run associated variants. Rather, we first identified regions with a high density of variants with large allele frequency differences between winter-run and non-winter-run fish in the CCV. Subsequently we identified SNPs within those regions with particularly large allele frequency differences. This approach was taken to avoid targeting single, isolated SNPs with large allele frequency differences that may have resulted merely from sampling variation, which was a concern because we had whole genome sequencing data from only 16 winter run fish.

More specifically, we calculated allele frequencies for 16 winter-run fish and 84 non-winter-run fish from the(Thompson *et al*., 2020) variant data VCF files using ANGSD version 0.921. Sites were retained if at least 62.5% of samples in each group had read data (10 of 16 for the winter run and 50 of 80 for the non-winter run), resulting in 7,295,001 SNPs for downstream analysis. We first investigated the distribution throughout the genome of | *d*|, the absolute difference of the alternate allele frequency between the two groups (Figure S8). This revealed that many loci had large values of |*d*|, but there were several regions in the genome, in particular, with prominent peaks in allele frequency difference and one or more SNPs apparently fixed for alternate alleles between the two sample groups.

To leverage information from multiple SNPs to identify regions in the genome with large allele frequency differences, we calculated the fraction of SNPs with |*d*| > 0.5 that also had |*d*| > 0.9 within non-overlapping 100 kb sliding windows throughout the genome. This metric indicated several prominent peaks. We focused on all of those 100 kb windows in which more than 12.4% of sites with |*d*| > 0.5 also had |*d*| > 0.9 which were also adjacent to at least one window in which > 10% of sites with |*d*| > 0.5 also had |*d*| > 0.9 (Figure S9). Within the windows found on four different chromosomes using the above criteria, we then attempted to design amplicons to type the SNPs at a subset of the sites within each window. We chose all sites with |*d*| > 0.975, as well as the 8 SNPs on each chromosome with the highest values of |*d*|, yielding 61 candidate SNPs (Figure S10).

Some of those 61 candidate SNPs were close enough that it was possible to consider amplifying them with PCR on 58 different short sequences. We used Primer3 (Untergasser *et al*., 2012) to return three possible primer-pair designs for each of the 58 sequences and then chose the primer pair with the fewest penalties, optimal target size, and most consistent melting temperatures. One primer pair was dropped from consideration because it amplified a large indel that was apparent in the sequence data which would have rendered the sequence too large to efficiently amplify and several others were dropped because the primers overlapped other amplicons. Finally, several amplicons with the lowest |*d*| on chromosome 16 (RefSeq NC_037112.1) were removed from consideration, leaving us with 48 amplicons to test for amplification and for evaluation of allele frequencies.

These 48 amplicons were amplified and sequenced in 192 fish—96 Feather River spring run and 96 winter run—on a MiSeq sequencer, and the variants were called using GATK. The resulting VCF file was used to estimate allele frequency differences between winter run and Feather River spring run. We also processed the sequence data using the R package, ‘microhaplot’ (https://github.com/ngthomas/microhaplot), and visually inspected loci for consistent allele depth ratio and numbers of haplotypes. We then chose 24 amplicons for further testing on the basis of allele frequency differences between Feather River spring run and winter run, number of haplotypes, and ease of scoring. These 24 markers were typed on a variety of fish over the course of a year, and we finally chose three to include in our California Chinook reference baseline: one amplicon on each of chromosomes 8, 12, and 16. The estimated frequencies of all the alleles present in the reference baseline in those three amplicons shows that there are not fixed differences at these markers between winter run and all other reporting units (Table S2).

**Figure S1.**
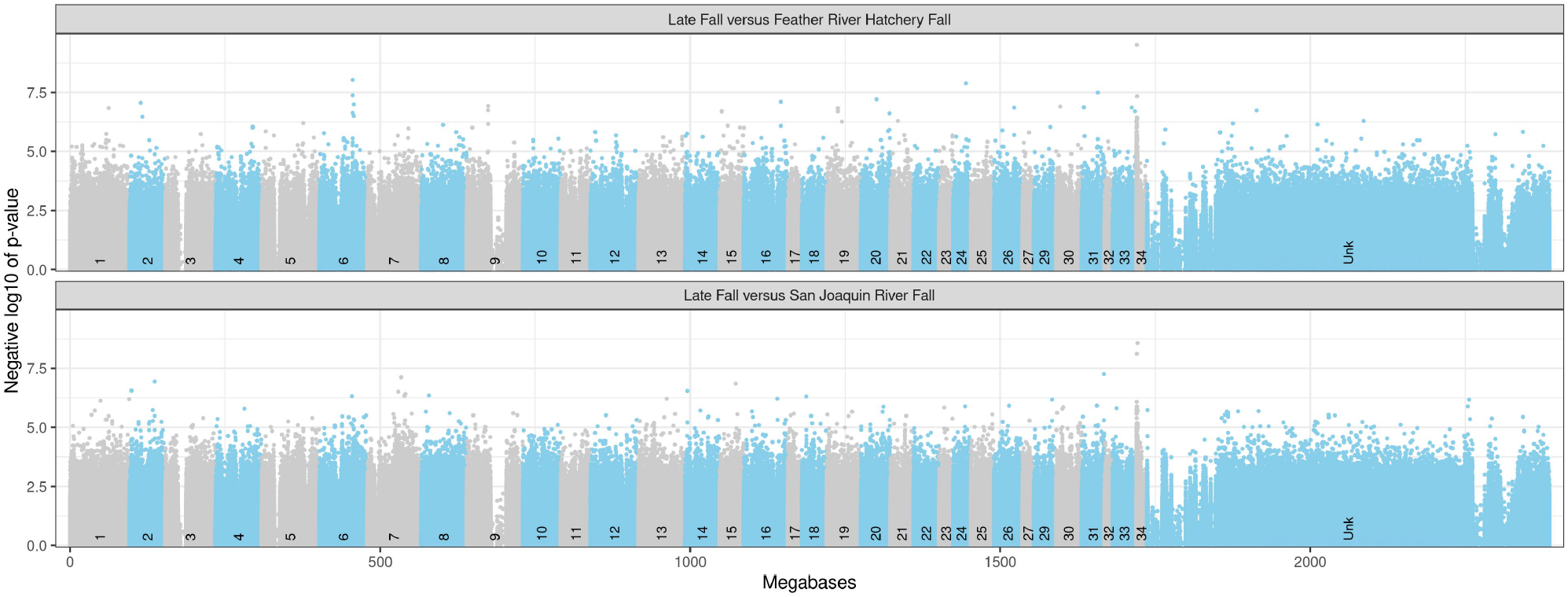
Negative log base 10 of association *p*-values for individual SNPs for late-fall versus fall run. *x* axis shows position in genome (in megabases), with color alternating by chromosome, as indicated by numbers above the *x*-axis. “Unk” refers to unplaced scaffolds in the Otsh_v1.0 genome assembly (Christensen *et al*., 2018). The upper panel is the comparison between late-fall and Feather River Hatchery fall, while the lower panel is the comparison of late-fall to San Joaquin River fall.

**Figure S2.**
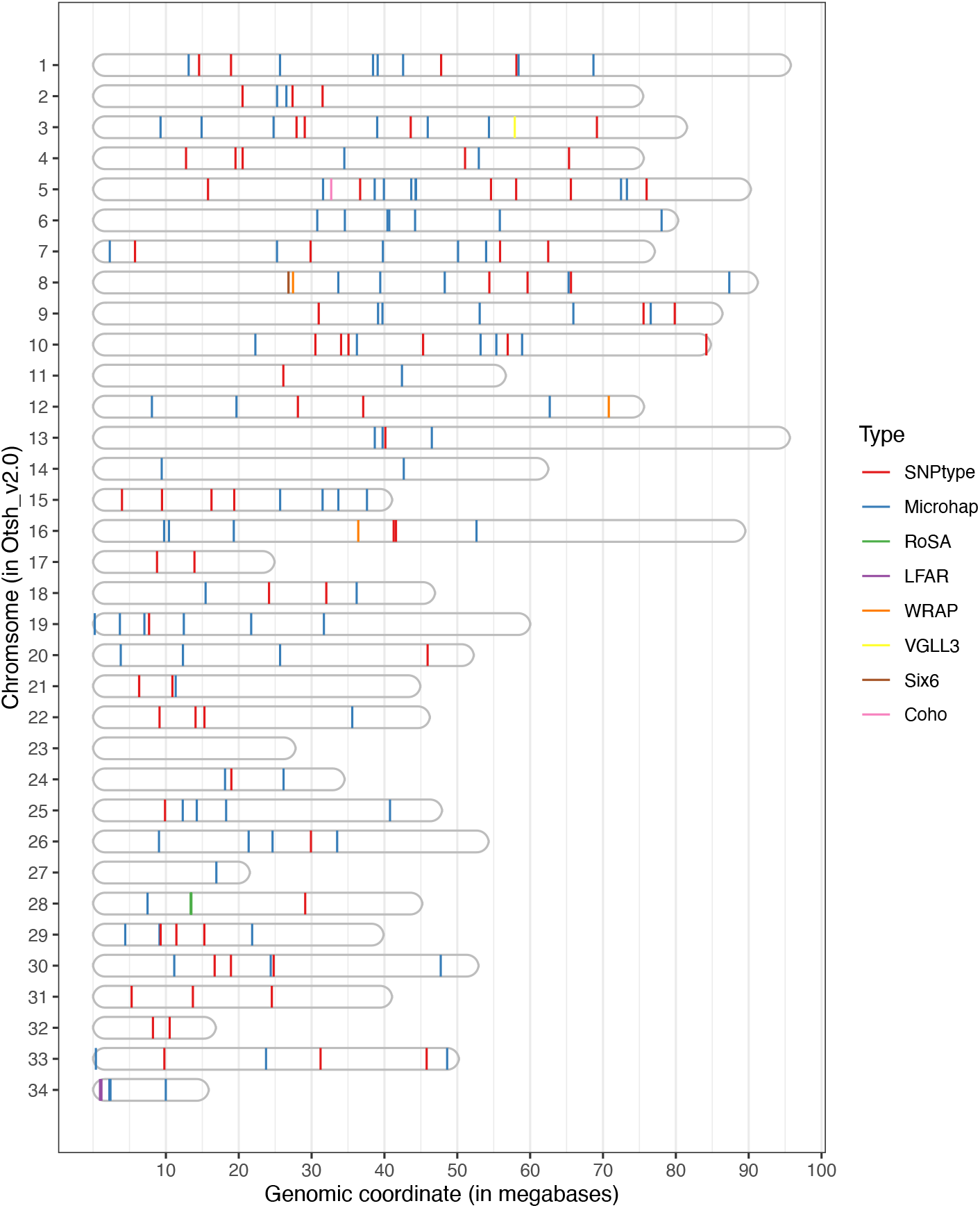
Genomic locations of amplicons in the Otsh_v2.0 assembly of the Chinook salmon genome. Color shows type of marker (see main text). The sex-ID marker is not included here because it aligns to a scaffold that is not part of a named chromosome/linkage group in the assembly.

**Figure S3.**
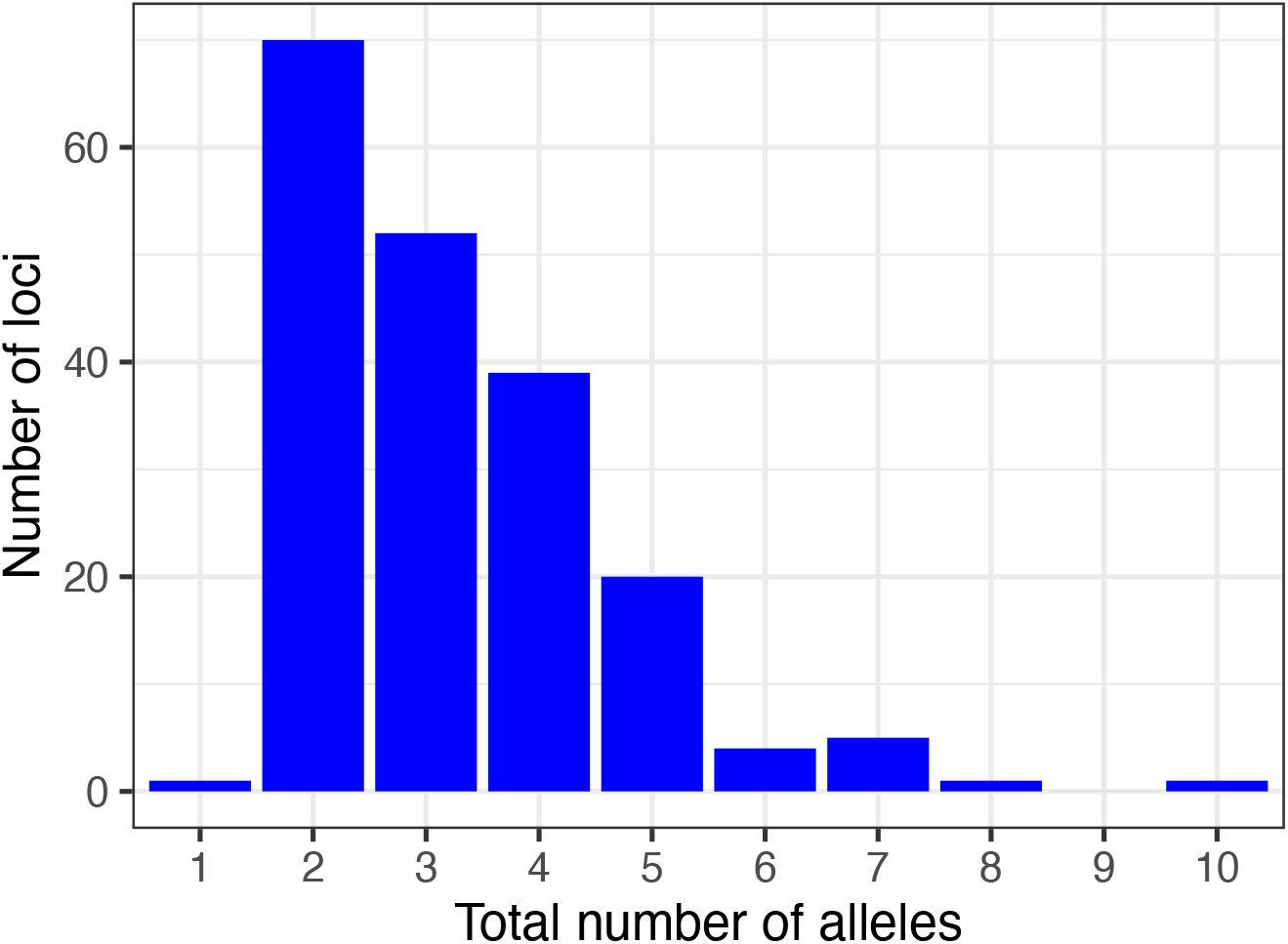
Number of loci with different total numbers of alleles in the data set. The one amplicon with only one allele is `Ots_coho001_05_32691399`, which is fixed for alternate alleles in Chinook vs. coho salmon. It is helpful in identifying coho samples that are misidentified during sampling as Chinook salmon.

**Figure S4.**
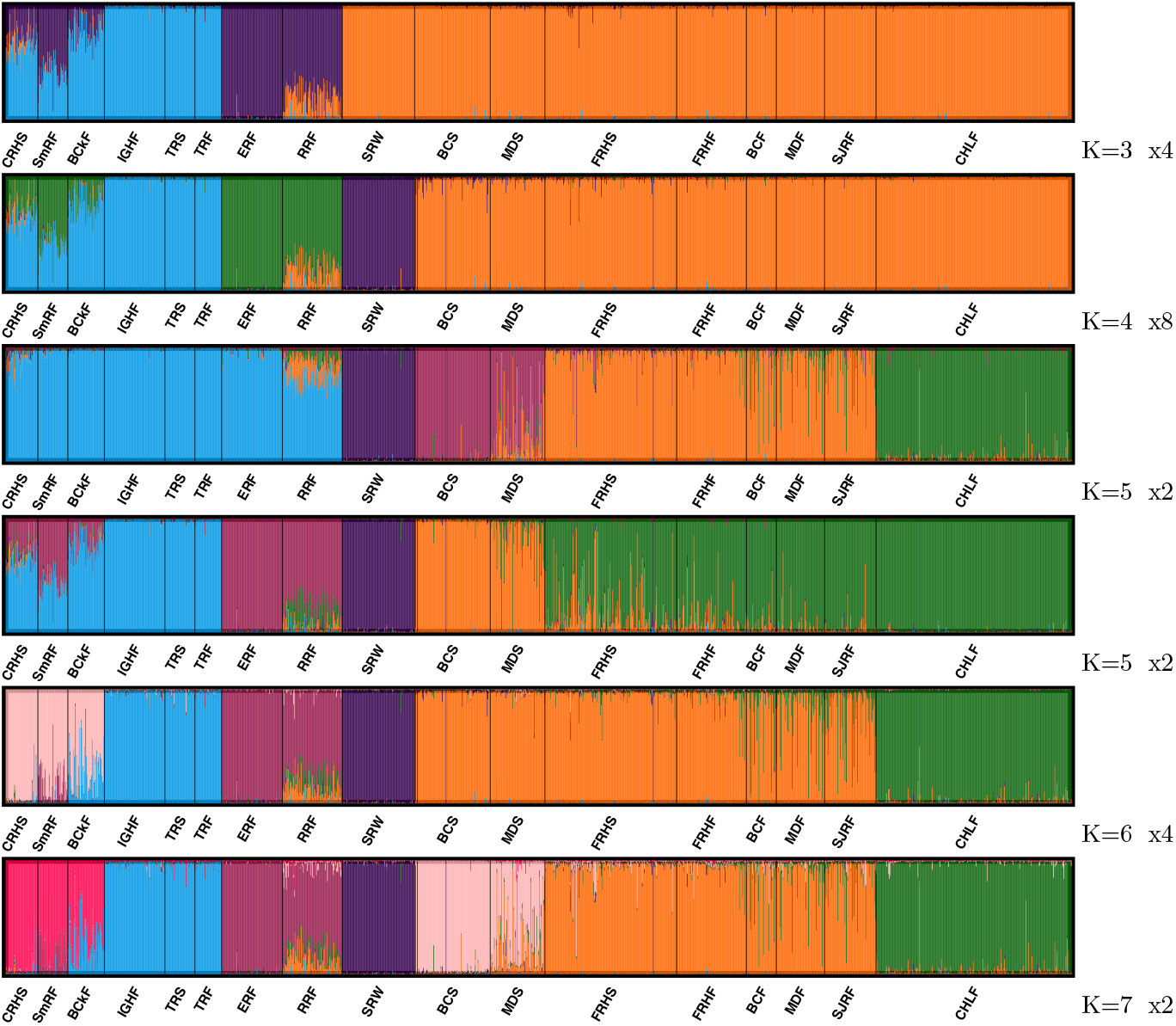
STRUCTURE minor modes found by CLUMPAK. At each value of *K* for which a minor mode was found, the plot is shown. *K* values and number of times each minor mode appeared out of 20 replicate runs of STRUCTURE appear to the lower right of each.

**Figure S5.**
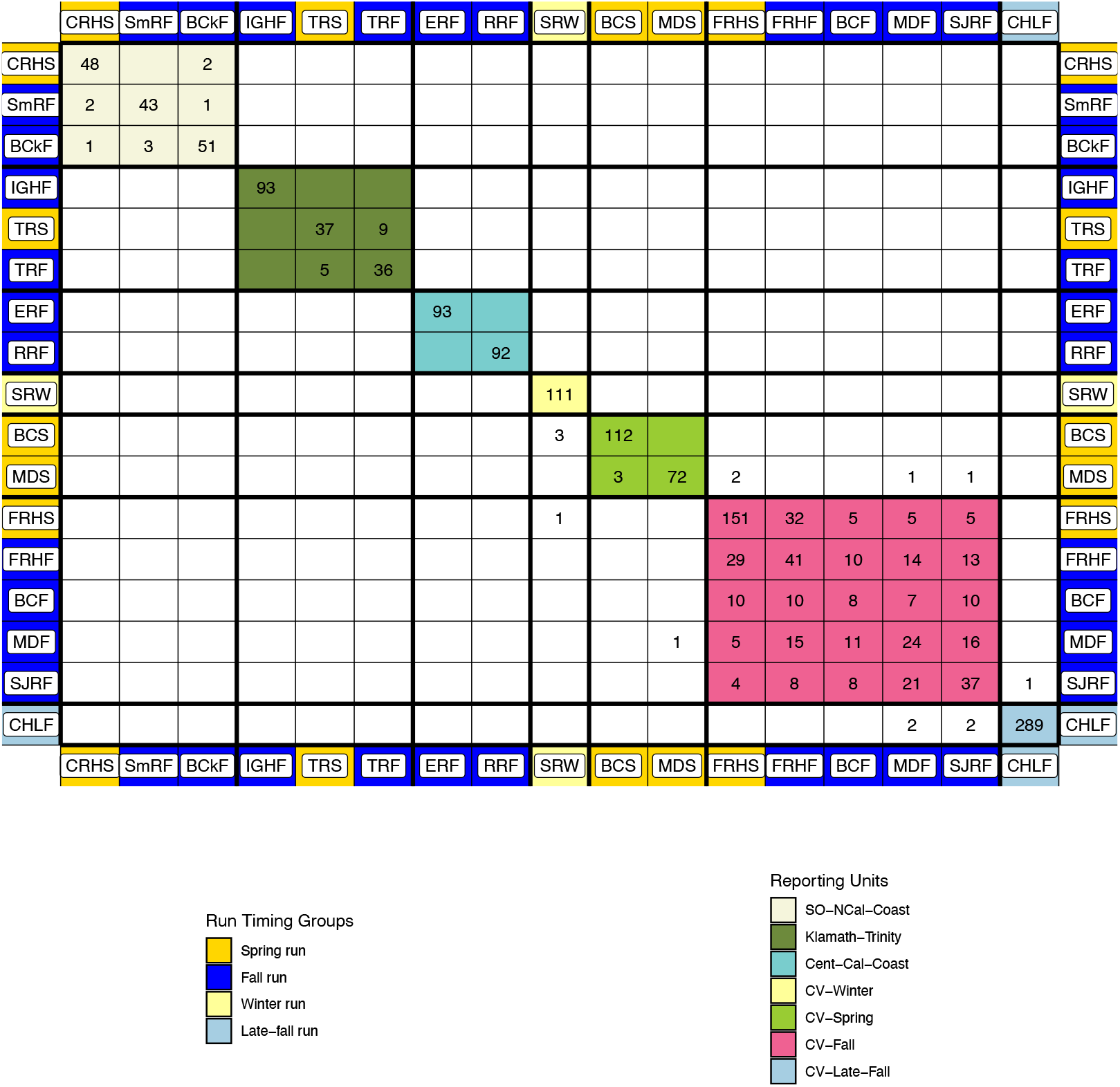
Assignment table like that in Figure 2b in the paper, but constrained so that only fish assigning to a reporting unit with scaled likelihood greater than 0.8 are included.

**Figure S6.**
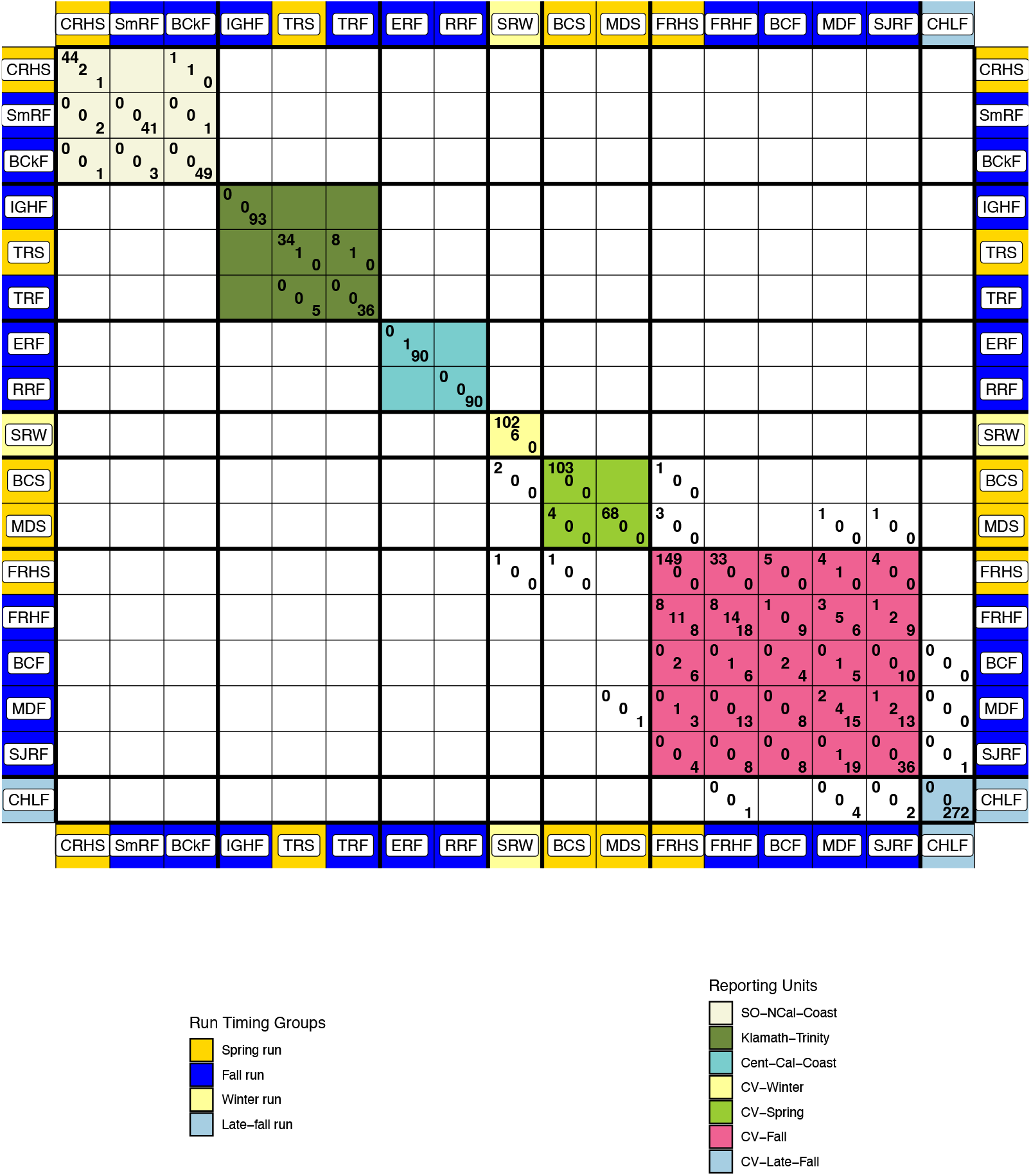
Assignment table like that in Figure 2b in the paper, but with numbers according to genotypes at the RoSA. In each cell, the top left entry gives the number of EE (early run allele homozygotes) genotypes, the middle entry is the number of EL genotypes (heterozygotes), and the bottom right entry gives the number of LL (late-run homozygote) genotypes.

**Figure S7.**
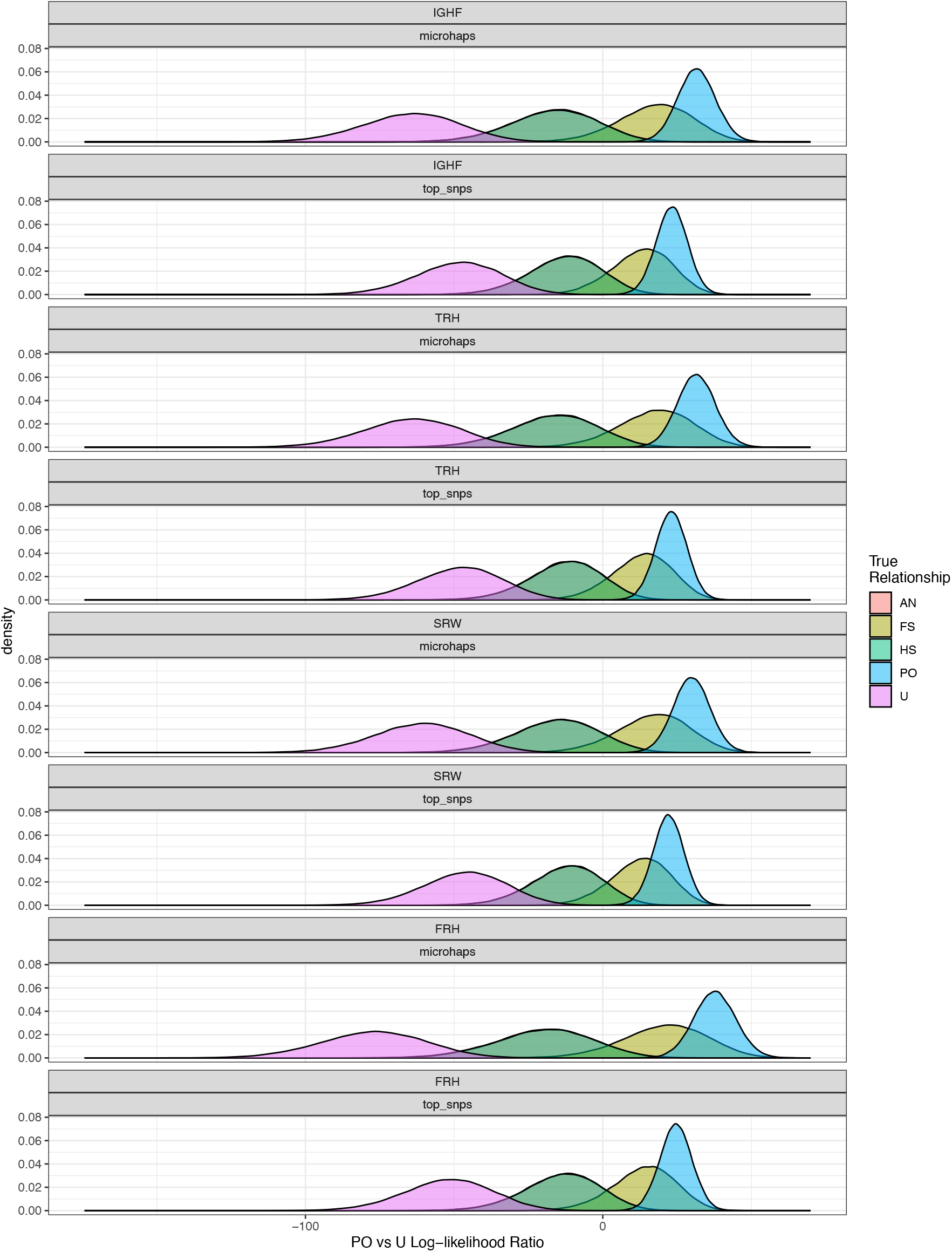
A comparison of the distribution of log-likelihood ratios when using all the alleles at each amplicon typed as microhaplotypes (“microhaps” in the panel headers) versus using just a single (the most informative) SNP from each amplicon (“top_snps” in the panel headers). Results shown for four hatchery collections in California (FRH: Feather River Hatchery [spring and fall]; IGH: Iron Gate Hatchery; SRW: Sacramento Winter Run; TRH: Trinity River Hatchery [spring and fall]). The density plots show the distribution of log-likelihood ratios for Parent-Offspring vs Unrelated for four different relationships: PO = parent offspring; FS = full-sibling; HS = Half sibling; AN = avuncular (i.e., aunt-niece, etc.) The distributions for AN and HS overlap completely. Note that the overlap between FS and PO occurs for the PO vs U likelihood ratio, but is nearly eliminated with the PO vs FS likelihood ratio, allowing these two relationships to be resolved accurately.

**Figure S8.**
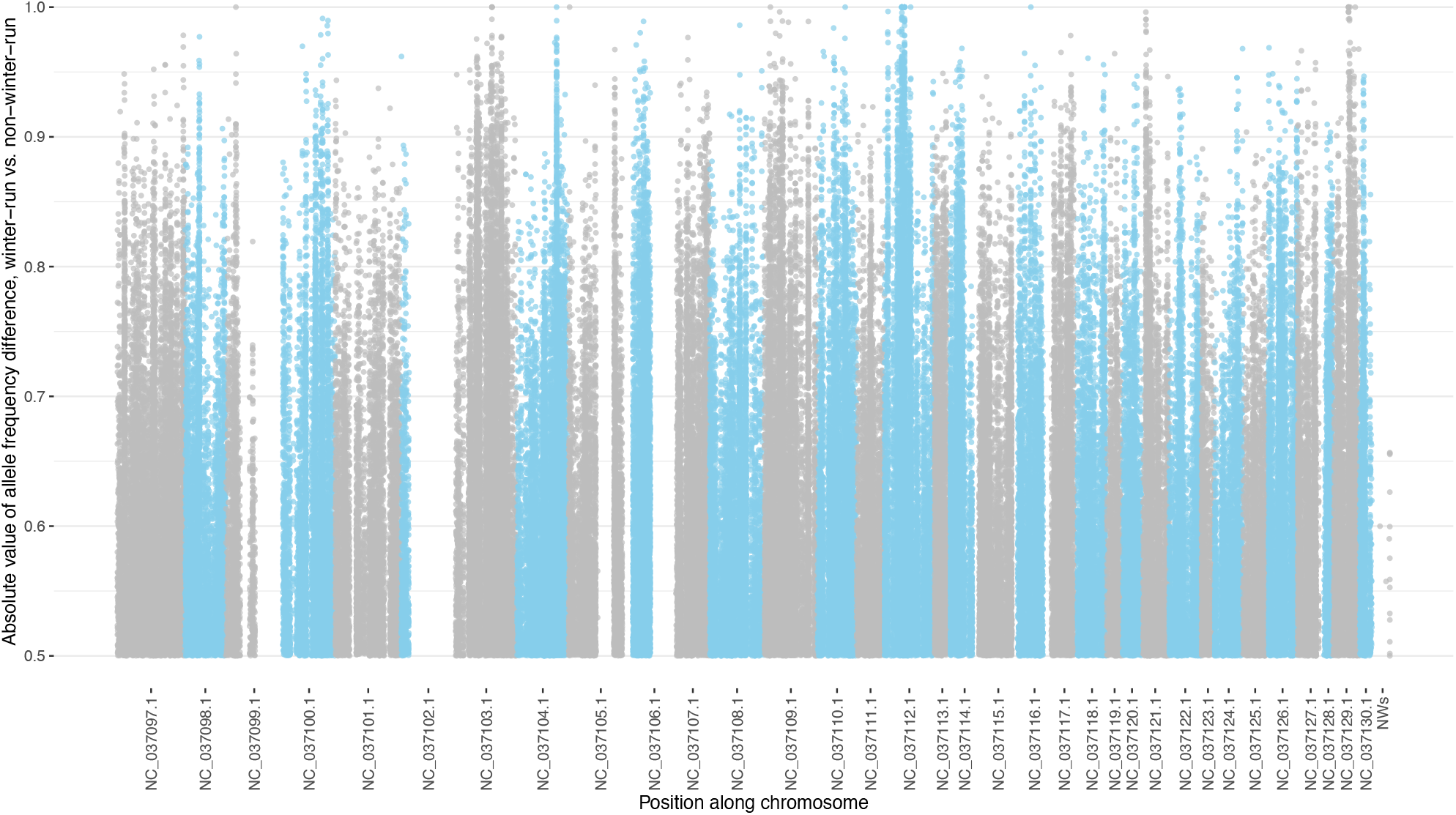
Absolute difference in allele frequency between winter run and non-winter run fish of the CCV. The plot shows only those SNPs with at least a frequency difference of 0.5 between the two groups. Each point is a SNP. The *x* axis shows position in the Otsh_v1.0 genome with color alternating by chromosome as indicated by RefSeq names on plot.

**Figure S9.**
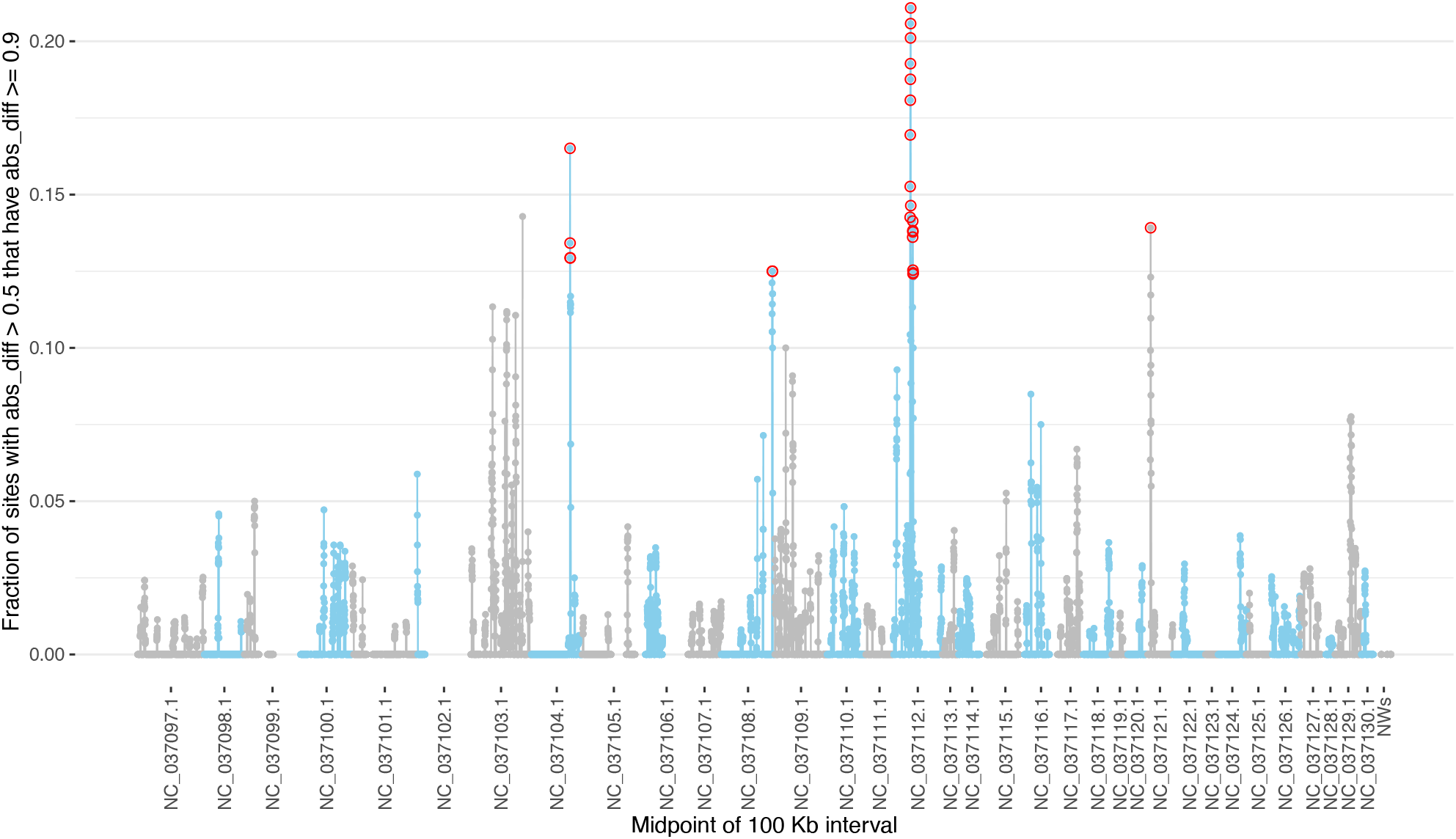
Values within 100 kb sliding windows of the fraction of sites with |*d*| (absolute difference in allele frequency between winter run and non-winter run fish of the CCV) greater than 0.5 that also have |*d*| > 0.9. The *x* axis shows position in the Otsh_v1.0 genome with color alternating by chromosome as indicated by RefSeq names on plot. Red circles denote sliding windows chosen for further investigation for candidate markers.

**Figure S10.**
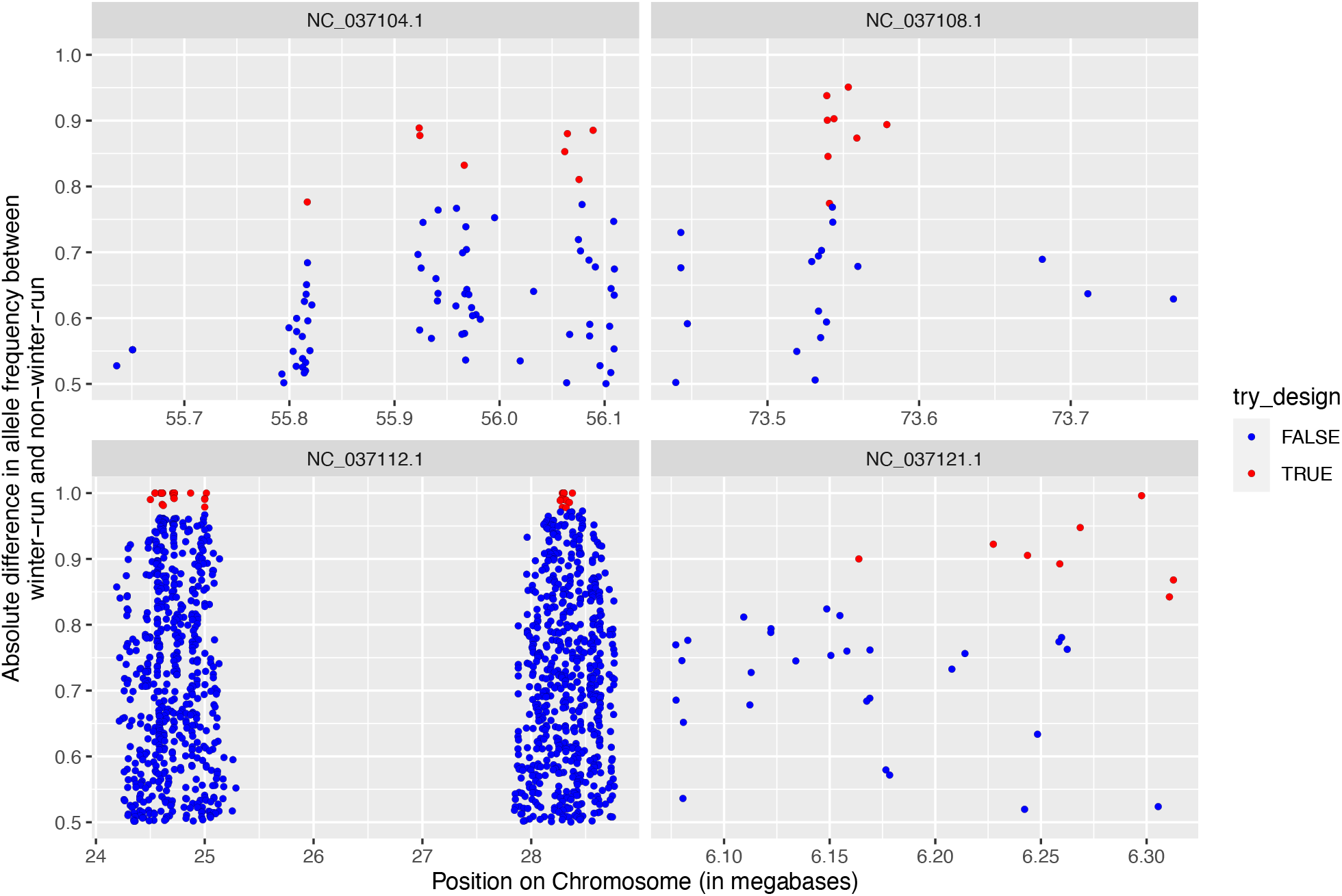
Genomic positions and values of |*d*| (absolute difference in allele frequency between winter run and non-winter run fish of the CCV) for the 61 candidate SNPs (in red) to design for winter-run-associated polymorphisms (WRAPs). Other sites are shown in blue. The *x*-axis shows position on each chromsome in the Otsh_v1.0 assembly. The chromsome name is indicated in the facet headers by RefSeq name

**Table S1.**
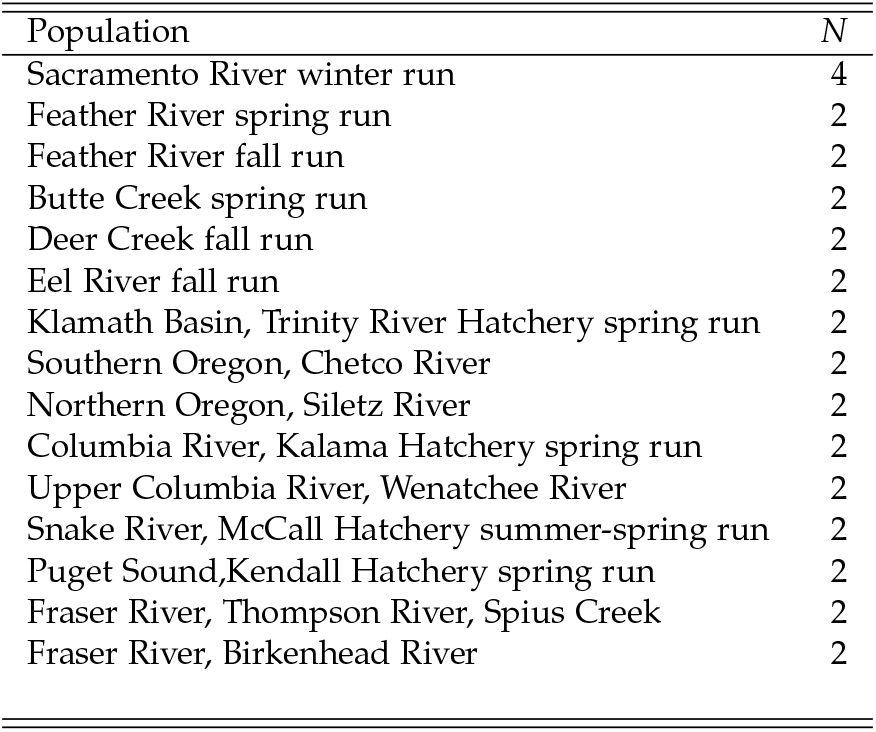
Numbers of fish from different collections/populations used in the microhaplotype discovery effort. *N* is the number of diploid individuals included in the ascertainment panel.

**Table S2.** Allele frequencies across reporting units of the three winter-run associated markers. Genome coordinates of SNPs are from Otsh_v2.0. Alleles are listed according to the SNP nucleotide at the variable SNPs within the amplicon.

| Locus | Allele | CV-Winter | CV-Spring | CV-Fall | CV-Late-Fall | Cent-Cal-Coast | Klamath-Trinity | SO-NCal-Coast |
| --- | --- | --- | --- | --- | --- | --- | --- | --- |
| Chr08:27450181 | C | 0.838 | 0.257 | 0.088 | 0.039 | 0.015 | 0.022 | 0.016 |
| Chr08:27450181 | A | 0.162 | 0.743 | 0.912 | 0.961 | 0.985 | 0.978 | 0.984 |
| Chr12:70794116 | GAA | 0.815 | 0.032 | 0.033 | 0.017 | 0.000 | 0.000 | 0.000 |
| Chr12:70794116 | GAT | 0.149 | 0.860 | 0.817 | 0.734 | 0.700 | 0.720 | 0.686 |
| Chr12:70794116 | AAT | 0.032 | 0.087 | 0.142 | 0.234 | 0.022 | 0.026 | 0.024 |
| Chr12:70794116 | GTT | 0.005 | 0.021 | 0.008 | 0.015 | 0.278 | 0.254 | 0.291 |
| Chr16:36409831 | CCG | 0.797 | 0.026 | 0.007 | 0.003 | 0.000 | 0.009 | 0.192 |
| Chr16:36409831 | TCG | 0.162 | 0.263 | 0.336 | 0.437 | 0.195 | 0.430 | 0.371 |
| Chr16:36409831 | TCA | 0.036 | 0.616 | 0.610 | 0.507 | 0.485 | 0.547 | 0.415 |
| Chr16:36409831 | TGG | 0.005 | 0.095 | 0.048 | 0.053 | 0.320 | 0.015 | 0.022 |

## References

Abadía-Cardoso A, Anderson EC, Pearse DE, Garza JC (2013) Large-scale parentage analysis reveals reproductive patterns and heritability of spawn timing in a hatchery population of steelhead (Oncorhynchus mykiss). Molecular Ecology, 22, 4733–4746.

Allendorf FW, Phelps SR (1981) Use of allelic frequencies to describe population structure. Canadian Journal of Fisheries and Aquatic Sciences, 38, 1507–1514.

Anderson E (2010) Assessing the power of informative subsets of loci for population assignment: standard methods are upwardly biased. Molecular ecology resources, 10, 701–710.

Anderson E, Thompson E (2002) A model-based method for identifying species hybrids using multilocus genetic data. Genetics, 160, 1217–1229. https://citeseerx.ist.psu.edu/document?repid=rep1&type=pdf&doi=67528b8c6ebf8afd79d9129344e489e1bbe238e2.

Anderson EC, Garza JC (2005) A description of full parental genotyping, Technical report, Unpublished report on file with the National Oceanic and Atmospheric Administration. Seattle, WA.

Anderson EC, Garza JC (2006) The power of single-nucleotide polymorphisms for large-scale parentage inference. Genetics, 172, 2567–2582.

Anderson EC, Waples RS, Kalinowski ST (2008) An improved method for predicting the accuracy of genetic stock identification. Canadian Journal of Fisheries and Aquatic Sciences, 65, 1475–1486.

Ayala FJ, Powell JR (1972) Allozymes as diagnostic characters of sibling species of Drosophila. Proceedings of the National Academy of Sciences, 69, 1094–1096.

Baetscher DS, Clemento AJ, Ng TC, Anderson EC, Garza JC (2018) Microhaplotypes provide increased power from short-read DNA sequences for relationship inference. Molecular Ecology Resources, 18, 296–305.

Barclay AW, Habicht C (2019) Genetic Baseline for Cook Inlet Coho Salmon and Evaluations for Mixed Stock Analysis, Technical report, Alaska Department of Fish and Game, Divisions of Sport Fish and Commercial Fisheries.

Barson NJ, Aykanat T, Hindar K, Baranski M, Bolstad GH, Fiske P, Jacq C, Jensen AJ, Johnston SE, Karlsson S, et al. (2015) Sex-dependent dominance at a single locus maintains variation in age at maturity in salmon. Nature, 528, 405–408.

Beacham TD, Lapointe M, Candy JR, Miller KM, Withler RE (2004) DNA in action: rapid application of DNA variation to sockeye salmon fisheries management. Conservation Genetics, 5, 411–416.

Beacham TD, Wallace CG, Jonsen K, McIntosh B, Candy JR, Horst K, Lynch C, Willis D, Luedke W, Kearey L, et al. (2021) Parentage-based tagging combined with genetic stock identification is a cost-effective and viable replacement for coded-wire tagging in large-scale assessments of marine Chinook salmon fisheries in British Columbia, Canada. Evolutionary Applications, 14, 1365–1389.

Bertho S, Herpin A, Jouanno E, Yano A, Bobe J, Parrinello H, Journot L, Guyomard R, Muller T, Swanson P, et al. (2022) A nonfunctional copy of the salmonid sex-determining gene (sdY) is responsible for the “apparent” XY females in Chinook salmon, Oncorhynchus tshawytscha. G3, 12, jkab451.

Beulke AK, Abadía-Cardoso A, Pearse DE, Goetz LC, Thompson NF, Anderson EC, Garza JC (2023) Distinct patterns of inheritance shape life-history traits in steelhead trout. Molecular Ecology, 32, 6896–6912.

Bjorkstedt EP, Spence BC, Garza JC, Hankin DG, Fuller D, Jones WE, Smith JJ, Macedo R (2005) An analysis of historical population structure for evolutionary significant units of Chinook salmon, coho salmon, and steelhead in the North-Central California Coast Recovery Domain, Technical report, NOAA Technical Memorandum.

Campbell NR, Harmon SA, Narum SR (2015) Genotyping-in-Thousands by sequencing (GT-seq): A cost effective SNP genotyping method based on custom amplicon sequencing. Molecular Ecology Resources, 15, 855–867.

Catchen J, Hohenlohe PA, Bassham S, Amores A, Cresko WA (2013) Stacks: an analysis tool set for population genomics. Molecular Ecology, 22, 3124–3140.

Cingolani P, Platts A, Coon M, Nguyen T, Wang L, Land S, Lu X, Ruden D (2012) A program for annotating and predicting the effects of single nucleotide polymorphisms, SnpEff: SNPs in the genome of Drosophila melanogaster strain w1118; iso-2; iso-3. Fly, 6, 80–92.

Clemento A, Abadía-Cardoso A, Starks H, Garza J (2011) Discovery and characterization of single nucleotide polymorphisms in Chinook salmon, Oncorhynchus tshawytscha. Molecular Ecology Resources, 11, 50–66.

Clemento AJ, Crandall ED, Garza JC, Anderson EC (2014) Evaluation of a single nucleotide polymorphism baseline for genetic stock identification of Chinook Salmon (Oncorhynchus tshawytscha) in the California Current large marine ecosystem. Fishery Bulletin, 112.

DeSaix MG, Rodriguez MD, Ruegg KC, Anderson EC (2024) Population assignment from genotype likelihoods for low-coverage whole-genome sequencing data. Methods in Ecology and Evolution, 15, 493–510.

Falush D, Stephens M, Pritchard JK (2003) Inference of population structure using multilocus genotype data: linked loci and correlated allele frequencies. Genetics, 164, 1567–1587.

Fisher FW (1994) Past and present status of Central Valley Chinook salmon. Conservation Biology, 8, 870–873.

Fry DH (1961) King salmon spawning stocks of the California Central Valley, 1940-1959. California Fish and Game, 47, 55–7l.

Garrison E, Marth G (2012) Haplotype-based variant detection from short-read sequencing. arXiv preprint arXiv:1207.3907.

Garza JC, Anderson E (2007) Large scale parentage inference as an alternative to coded-wire tags for salmon fishery management, in: PSC Genetic Stock Identification Workshop (May and September 2007): Logistics Workgroup final report and recommendations. Vancouver, British Columbia, Canada. Available at http://www.psc.org/info_genetic_stock_id.htm#REPORTS, Pacific Salmon Commission.

Gilbey J, Coughlan J, Wennevik V, Prodöhl P, Stevens JR, Garcia de Leaniz C, Ensing D, Cauwelier E, Cherbonnel C, Consuegra S, et al. (2018) A microsatellite baseline for genetic stock identification of European Atlantic salmon (Salmo salar L.). ICES Journal of Marine Science, 75, 662–674.

Goudet J, Jombart T (2022) hierfstat: Estimation and Tests of Hierarchical F-Statistics. R package version 0. 5-11.

Hardy GH (1908) Mendelian proportions in a mixed population. Science, 28, 49–50.

Hasselman DJ, Anderson EC, Argo EE, Bethoney ND, Gephard SR, Post DM, Schondelmeier BP, Schultz TF, Willis TV, Palkovacs EP (2016) Genetic stock composition of marine bycatch reveals disproportional impacts on depleted river herring genetic stocks. Canadian Journal of Fisheries and Aquatic Sciences, 73, 951–963.

Healey M (1991) Life history of Chinook salmon (Oncorhynchus tshawytscha), in: Pacific salmon life histories, 311–394, University of British Columbia Press Vancouver.

Hedrick PW, Hedgecock D (1994) Effective population size in winter-run Chinook salmon. Conservation Biology, 8, 890–892.

Hendricks S, Anderson EC, Antao T, Bernatchez L, Forester BR, Garner B, Hand BK, Hohenlohe PA, Kardos M, Koop B, et al. (2018) Recent advances in conservation and population genomics data analysis. Evolutionary Applications, 11, 1197–1211.

Hess JE, Whiteaker JM, Fryer JK, Narum SR (2014) Monitoring stock-specific abundance, run timing, and straying of Chinook salmon in the Columbia River using genetic stock identification (GSI). North American Journal of Fisheries Management, 34, 184–201.

Hillary R, Bravington M, Patterson T, Grewe P, Bradford R, Feutry P, Gunasekera R, Peddemors V, Werry J, Francis M, et al. (2018) Genetic relatedness reveals total population size of white sharks in eastern Australia and New Zealand. Scientific reports, 8, 2661.

Horn RL, Hess M, Harmon S, Hess J, Delomas TA, Campbell MR, Narum S (2023) Multigeneration pedigrees to monitor hatchery broodstock composition and genetic variation of spring/summer Chinook almon in the Columbia River basin. North American Journal of Fisheries Management, 43, 794–820.

Kent WJ (2002) BLAT—the BLAST-like alignment tool. Genome Research, 12, 656–664.

Kim SY, Lohmueller KE, Albrechtsen A, Li Y, Korneliussen T, Tian G, Grarup N, Jiang T, Andersen G, Witte D, Jorgensen T, Hansen T, Pedersen O, Wang J, Nielsen R (2011) Estimation of allele frequency and association mapping using next-generation sequencing data. BMC Bioinformatics, 12, 231.

Kinziger AP, Hellmair M, Hankin DG, Garza JC (2013) Contemporary population structure in Klamath River basin Chinook salmon revealed by analysis of microsatellite genetic data. Transactions of the American Fisheries Society, 142, 1347–1357.

Kopelman NM, Mayzel J, Jakobsson M, Rosenberg NA, Mayrose I (2015) Clumpak: a program for identifying clustering modes and packaging population structure inferences across K. Molecular Ecology Resources, 15, 1179–1191.

Koressaar T, Remm M (2007) Enhancements and modifications of primer design program Primer3. Bioinformatics, 23, 1289–1291.

Korneliussen TS, Albrechtsen A, Nielsen R (2014) ANGSD: Analysis of next generation sequencing data. BMC Bioinformatics, 15, 356.

Lange K, Papp JC, Sinsheimer JS, Sripracha R, Zhou H, Sobel EM (2013) Mendel: the Swiss army knife of genetic analysis programs. Bioinformatics, 29, 1568–1570.

Li H, Durbin R (2009) Fast and accurate short read alignment with Burrows–Wheeler transform. Bioinformatics, 25, 1754–1760.

Li H, Handsaker B, Wysoker A, Fennell T, Ruan J, Homer N, Marth G, Abecasis G, Durbin R, Subgroup GPDP (2009) The sequence alignment/map format and SAMtools. Bioinformatics, 25, 2078–2079.

Lindley ST, Grimes CB, Mohr MS, Peterson WT, Stein JE, Anderson JJ, Botsford LW, Bottom DL, Busack CA, Collier TK, et al. (2009) What caused the Sacramento River fall Chinook stock collapse, Technical Memorandum NOAA-TM-NMFS-SWFSC 447, NOAA.

Magoč T, Salzberg SL (2011) FLASH: fast length adjustment of short reads to improve genome assemblies. Bioinformatics, 27, 2957–2963.

McKinney GJ, Seeb JE, Seeb LW (2017) Managing mixed-stock fisheries: genotyping multi-SNP haplotypes increases power for genetic stock identification. Canadian Journal of Fisheries and Aquatic Sciences, 74, 429–434.

Meek MH, Baerwald MR, Stephens MR, Goodbla A, Miller MR, Tomalty KM, May B (2016) Sequencing improves our ability to study threatened migratory species: Genetic population assignment in California’s Central Valley Chinook salmon. Ecology and Evolution, 6, 7706–7716.

Meek MH, Stephens MR, Goodbla A, May B, Baerwald MR (2020) Identifying hidden biocomplexity and genomic diversity in Chinook salmon, an imperiled species with a history of anthropogenic influence. Canadian Journal of Fisheries and Aquatic Sciences, 77, 534–547.

Milner GB, Teel DJ, Utter FM (1982) Gentic Identification Study Annual Progress Report, FY 81, Technical report, Coastal Zone and Estuarine Studies Division Northwest and Alaska Fisheries Center National Marine Fisheries Service National Oceanic and Atmospheric Administration.

Milner GB, Teel DJ, Utter FM, Winans GA (1985) A genetic method of stock identification in mixed populations of Pacific salmon, Oncorhynchus spp. Marine Fisheries Review, 47, 1–8.

Moran BM, Anderson EC (2019) Bayesian inference from the conditional genetic stock identification model. Canadian Journal of Fisheries and Aquatic Sciences, 76, 551–560.

Myers JM, Kope RG, Bryant GJ, Teel D, Lierheimer LJ, Wainwright TC, Grant WS, Waknitz FW, Neely K, Lindley ST, Waples RS (1998) Status review of Chinook salmon from Washington, Idaho, Oregon, and California, Technical report, U.S. Dept. of Commerce, National Oceanic and Atmospheric Administration, National Marine Fisheries Service, Seattle, WA.

Nandor GF, Longwill JR, Webb DL (2010) Overview of the coded wire tag program in the greater Pacific region of North America. PNAMP Special Publication: Tagging, Telemetry and Marking Measures for Monitoring Fish Populations—A compendium of new and recent science for use in informing technique and decision modalities: Pacific Northwest Aquatic Monitoring Partnership Special Publication, 2, 5–46.

Pella J, Masuda M (2006) The Gibbs and split merge sampler for population mixture analysis from genetic data with incomplete baselines. Canadian Journal of Fisheries and Aquatic Sciences, 63, 576–596.

Peterson BK, Weber JN, Kay EH, Fisher HS, Hoekstra HE (2012) Double digest RADseq: an inexpensive method for de novo SNP discovery and genotyping in model and non-model species. PloS one, 7, e37135.

Pritchard JK, Stephens M, Donnelly P (2000) Inference of population structure using multilocus genotype data. Genetics, 155, 945–959.

R Core Team (2024) R: A Language and Environment for Statistical Computing, R Foundation for Statistical Computing, Vienna, Austria.

Satterthwaite WH, Ciancio J, Crandall E, Palmer-Zwahlen ML, Grover AM, O’Farrell MR, Anderson EC, Mohr MS, Garza JC (2015) Stock composition and ocean spatial distribution inference from California recreational Chinook salmon fisheries using genetic stock identification. Fisheries Research, 170, 166–178.

Seeb L, Antonovich A, Banks MA, Beacham T, Bellinger M, Blankenship S, Campbell M, Decovich N, Garza J, Guthrie Iii C, et al. (2007) Development of a standardized DNA database for Chinook salmon. Fisheries, 32, 540–552.

Smouse PE, Waples RS, Tworek JA (1990) A genetic mixture analysis for use with incomplete source population data. Canadian Journal of Fisheries and Aquatic Sciences, 47, 620–634.

Steele CA, Anderson EC, Ackerman MW, Hess MA, Campbell NR, Narum SR, Campbell MR (2013) A validation of parentage-based tagging using hatchery steelhead in the Snake River basin. Canadian Journal of Fisheries and Aquatic Sciences, 70, 1046–1054.

Steele CA, Hess M, Narum S, Campbell M (2019) Parentage-based tagging: Reviewing the implementation of a new tool for an old problem. Fisheries, 44, 412–422.

Thompson NF, Anderson EC, Clemento AJ, Campbell MA, Pearse DE, Hearsey JW, Kinziger AP, Garza JC (2020) A complex phenotype in salmon controlled by a simple change in migratory timing. Science, 370, 609–613.

Thompson TQ, Bellinger MR, O’Rourke SM, Prince DJ, Stevenson AE, Rodrigues AT, Sloat MR, Speller CF, Yang DY, Butler VL, et al. (2019) Anthropogenic habitat alteration leads to rapid loss of adaptive variation and restoration potential in wild salmon populations. Proceedings of the National Academy of Sciences, 116, 177–186.

Thompson TQ, O’Leary S, O’Rourke S, Tarsa C, Baerwald MR, Goertler P, Meek MH (2024) Genomics and 20 years of sampling reveal phenotypic differences between subpopulations of outmigrating Central Valley Chinook salmon. Evolutionary Applications, 17, e13705.

Untergasser A, Cutcutache I, Koressaar T, Ye J, Faircloth BC, Remm M, Rozen SG (2012) Primer3—new capabilities and interfaces. Nucleic Acids Research, 40, e115–e115.

Urawa S, Sato S, Crane PA, Agler B, Josephson R, Azumaya T (2009) Stock-specific ocean distribution and migration of chum salmon in the Bering Sea and North Pacific Ocean. North Pacific Anadromous Fish Commission Bulletins, 5, 131–146.

Von Bargen J, Smith CT, Rueth J (2015) Development of a Chinook salmon sex identification SNP assay based on the growth hormone pseudogene. Journal of Fish and Wildlife Management, 6, 213–219.

Wang J (2004) Sibship reconstruction from genetic data with typing errors. Genetics, 166, 1963–1979.

Wang J (2023) Estimating current effective sizes of large populations from a single sample of genomic marker data: A comparison of estimators by simulations. Population Ecology.

Waples RS (1991) Pacific salmon, Oncorhynchus spp., and the definition of “species” under the Endangered Species Act. Marine Fisheries Review, 53, 11–22.

Waples RS, Waples RK (2011) Inbreeding effective population size and parentage analysis without parents. Molecular Ecology Resources, 11, 162–171.

Waters CD, Clemento A, Aykanat T, Garza JC, Naish KA, Narum S, Primmer CR (2021) Heterogeneous genetic basis of age at maturity in salmonid fishes. Molecular Ecology, 30, 1435–1456.

Weir BS, Cockerham CC (1984) Estimating F-statistics for the analysis of population structure. Evolution, 1358–1370.

Wilmot RL, Kondzela CM, Guthrie III CM, Moles A, Martinson E, Helle JH (1999) Origins of sockeye and chum salmon seized from the Chinese vessel Ying Fa, Technical report, (NPAFC Doc.) Auke Bay Fisheries Laboratory, Alaska Fisheries Science Center, NMFS, NOAA, 11305 Glacier Highway, Juneau, AK 99801-8626. 20 pp.

Yoshiyama RM, Fisher FW, Moyle PB (1998) Historical abundance and decline of Chinook salmon in the Central Valley region of California. North American Journal of Fisheries Management, 18, 487–521.

## References

Christensen KA, Leong JS, Sakhrani D, Biagi CA, Minkley DR, Withler RE, Rondeau EB, Koop BF, Devlin RH (2018) Chinook salmon (Oncorhynchus tshawytscha) genome and transcriptome. PloS one, 13, e0195461.

